# Aging affects reprogramming of murine pulmonary capillary endothelial cells after lung injury

**DOI:** 10.1101/2023.07.11.548522

**Authors:** Marin Truchi, Marine Gautier-Isola, Grégoire Savary, Hugo Cadis, Célia Scribe, Alberto Baeri, Arun Lingampally, Virginie Magnone, Cédric Girard-Riboulleau, Marie-Jeanne Arguel, Clémentine de Schutter, Julien Fassy, Nihad Boukrout, Romain Larrue, Nathalie Martin, Roger Rezzonico, Olivier Pluquet, Michael Perrais, Véronique Hofman, Charles-Hugo Marquette, Paul Hofman, Andreas Günther, Nicolas Ricard, Pascal Barbry, Sylvie Leroy, Kevin Lebrigand, Saverio Bellusci, Christelle Cauffiez, Georges Vassaux, Nicolas Pottier, Bernard Mari

## Abstract

Aging increases the risk of developing fibrotic diseases by hampering tissue regeneration after injury. Using longitudinal single-cell RNA-seq and spatial transcriptomics, here we compare the transcriptome of bleomycin-induced fibrotic lungs of young and aged mice, at 3 time points corresponding to the peak of fibrosis, regeneration and resolution. We find that lung injury shifts the transcriptomic profiles of three pulmonary capillary endothelial cells (PCEC) subpopulations. The associated signatures are linked to pro-angiogenic signaling with strong *Lrg1* expression and do not progress similarly throughout the resolution process between young and old animals. Moreover, part of this set of resolution-associated markers is also detected in PCEC from samples of patients with idiopathic pulmonary fibrosis. Finally, we find that aging also alters the transcriptome of PCEC which display typical pro-fibrotic and pro-inflammatory features. We propose that age-associated alterations in specific PCEC subpopulations may interfere with the process of lung progenitor differentiation, thus contributing to the persistent fibrotic process typical of human pathology.

## INTRODUCTION

Gas exchanges between the external environment and the cardiovascular system take place in the lung alveoli through the thin membrane separating alveolar epithelial cells (AEC) and the capillary plexus formed by pulmonary capillary endothelial cells (PCEC), supported by mesenchymal cells. This alveolar niche, whose homeostasis is constantly threatened by external aggressions, deploys high regeneration abilities through various mechanisms^1^. Inadequate responses can lead to the development of chronic diseases such as idiopathic pulmonary fibrosis (IPF), an age-related pathology characterized by the accumulation of collagen-producing mesenchymal cells (myofibroblasts) resulting in the remodeling of the alveolar environment and accumulation of extracellular matrix (ECM), leading to progressive destruction of the parenchyma^2^. IPF is the prototypical progressive interstitial lung disease (ILD) associated with progressive decline in lung function and a median survival time of less than five years after diagnosis. Although two treatments are available to manage the disease (ie: nintedanib and perfenidone) and several new therapeutic options are under investigation, there is no curative treatment for IPF to date^3^. The pathogenesis of IPF is complex and largely unknown. Current hypotheses suggest that repetitive microinjuries of unknown origin to AEC in the aging lung results in ineffective repair with excessive wound healing, chronic inflammation, apoptosis of AEC and endothelial cells (EC), activation of fibrogenic effector cells, formation of myofibroblasts foci, and finally excessive deposition of ECM resulting in the destruction of the lung architecture and the loss of lung functions^4^. Mechanisms linking aging to the development of pulmonary fibrosis are not fully understood and involve numerous cell-intrinsic and cell-extrinsic alterations. Among them, it has been proposed that genetic and epigenetic changes, loss of proteostasis, cellular senescence, metabolic dysfunction as well as stem cell exhaustion may contribute to the initiation and progression of fibrosis^5–7^.

Following lung injury, the restoration of parenchymal tissue integrity depends in particular on the proliferative capacities of PCEC^8^. Recent studies showed that, in physiological conditions, PCEC are functionally heterogeneous, with two distinct cell types conserved in multiple mammal species^9^. Aerocytes (aCap), are specialized in gas exchanges with type 1 AEC (AEC1), whereas general capillaries (gCap) act as progenitor-like cells able to self-renew and to differentiate into aCap to maintain the homeostasis of the microvascular endothelium. Multiple studies also report various EC subpopulations emerging after lung injury in mouse models^10–12^ or in the context of IPF^13^. In particular, a population of peri-bronchial venous ECs expressing COL15A1, and referred as “systemic venous” (SV EC), have been shown to expand in areas of bronchiolization and fibrosis in IPF patient while this EC subtype was restricted to the bronchial vasculature in healthy lungs^13, 14^. A recent study^15^ further characterized these COL15A1^pos^ systemic ECs in lungs from lethal COVID-19 and IPF patients and identified a second subpopulation lacking expression of venous marker genes while expressing markers commonly detected in gCap, referred as systemic capillary ECs (sCap). Nevertheless, the precise contribution of the physiological state-associated cell types in the formation of these pathological subpopulations is not fully known. Moreover, as the regenerative functions of EC decline with aging^16, 17^, the impact of aging on the dynamic of EC subpopulations associated with alveolar niche regeneration after injury-induced fibrosis remains largely unaddressed^18^.

The murine single intratracheal bleomycin instillation model is considered as the best-characterized animal model available for the preclinical assessment of potential therapies for pulmonary fibrosis^19^. After the acute inflammatory phase has subsided, bleomycin-instilled mice display AEC remodeling and fibrosis, but unlike patients with IPF, fibrosis progressively resolves, allowing to study the cellular and molecular mechanisms associated with alveolar repair and regeneration. Furthermore, studies assessing the response of older mice to bleomycin have revealed a more pronounced fibrosis, with mechanisms more reminiscent of IPF^20–22^. It has been notably suggested that fibrosis resolution or tissue regeneration is impaired in aged mice^23–26^ indicating that age-associated processes contribute to the persistent fibrosis that is observed in IPF.

In this context, the aim of the present study was to compare the fate of different cell populations present in the lungs after bleomycin-induced injury in young and aged animals, from the fibrotic response peak (14 days) until partial (28 days) and complete (60 days) resolution.

## RESULTS

### Aging is associated with a delay in the resolution of lung fibrosis

Lungs of young (7 weeks) and old (18 months) mice were collected 14, 28 or 60 days after bleomycin challenge (**Fig. 1A**). We noticed a shift of fibrosis resolution between young and aged mice, as visualized by total hydroxyproline assay (**Fig. 1B**), histological analyses (**Fig. 1C**) and Sirius red assay (**Supplemental Fig. S1A**) with a peak of fibrosis at day 14 in young mice followed by a progressive reduction of fibrosis at D28 while a strong fibrotic pattern was still detected at D28 in old mice. In both young and old animals, fibrosis resolution was almost complete at D60, with only a few limited fibrotic areas. We then performed a spatial transcriptomics experiment with the 10x Genomics Visium platform on lung sections from young or old mice taken 14 or 28 days after bleomycin instillation to compare the fibrotic and resolution stages between the two age groups (**Fig. 1D**). After filtering the genes and the low-quality spots, we processed the data, corresponding to a corpus of 2479 genes × 10813 spots count matrix and the associated spatial coordinates, using a deconvolution tool leveraging only on spatial gene expression profiles^27^. We identified 14 unique topics, *i.e.* modules of covarying genes proportionally represented in each spot and annotated them according to their differentially expressed genes (**Supplemental Fig. S1B-C and Supplementary table S1**).

**Figure 1.**
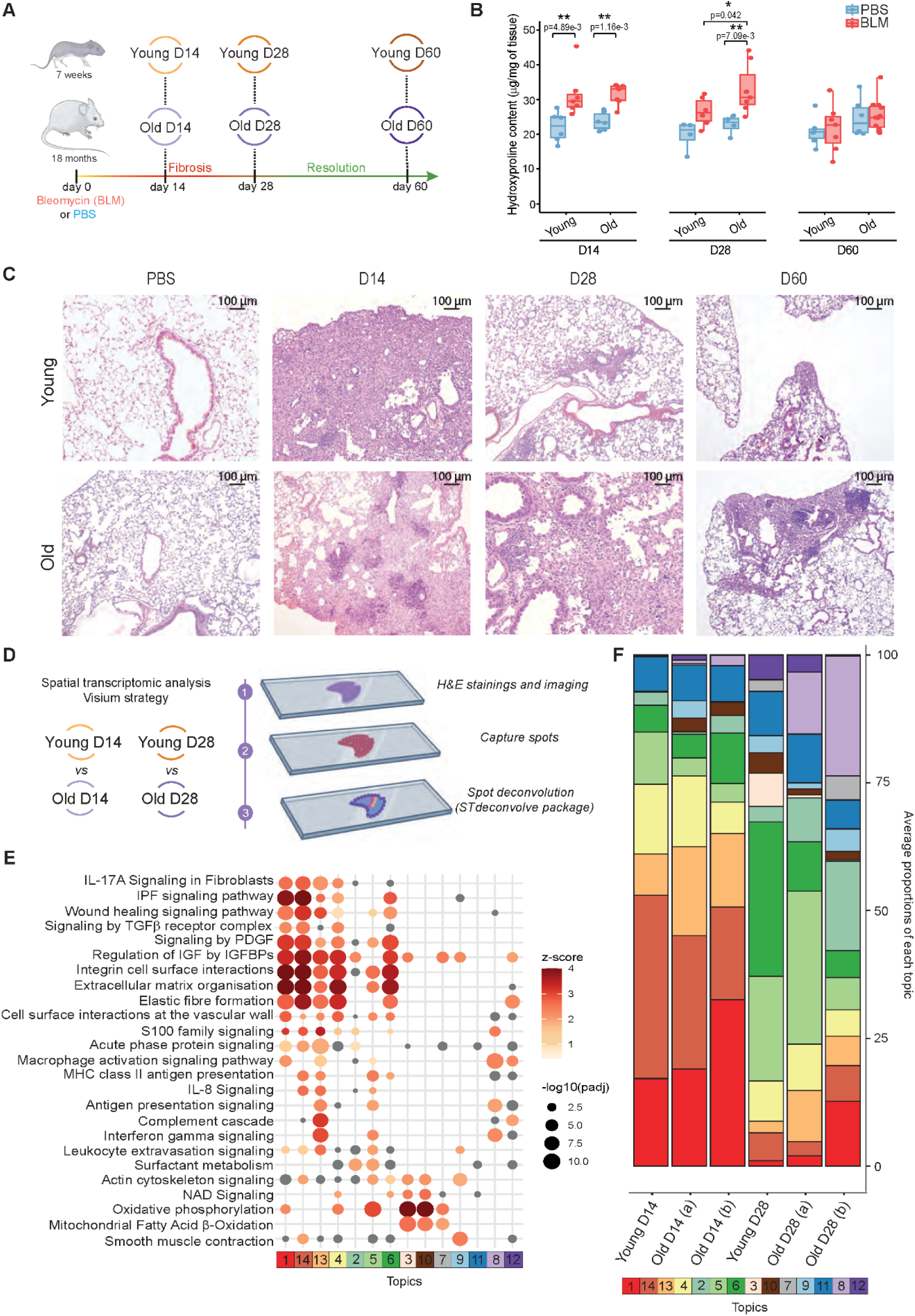
Analyses of fibrotic parameters in bleomycin-injured lungs from young and old mice. **(A)** Experimental design. Young (7 weeks) or old (18 months) mice were treated either with bleomycin (BLM) or PBS. Their lungs were collected at the bleomycin-induced fibrotic peak at day 14 and during lung regeneration (day 28) and fibrosis resolution (day 60). (**B)** Hydroxyproline quantification on BLM or PBS treated young or old mouse samples at indicated time points (n=4-10/group). * P<0.05. ** P<0.01. P values were calculated by two-way ANOVA test followed by a multiple comparisons test with Holm-Šídák correction. **(C)** Histological sections of control and fibrotic lungs from young and old mice using H&E at indicated time points (representative images, n=3). (**D)** Experimental design of spatial transcriptomics data from histological sections (H&E) of lungs from young (n=1) and old (n=2) mice challenged with bleomycin and collected at fibrotic peak (day 14, D14) and during regeneration (day 28, D28). Spots captured corresponding to tissue area were deconvoluted through ST deconvolve package in topics (n=14). (**E)** Functional annotations of topics by Ingenuity pathway. (**F)** Average proportion of each topic across deconvoluted spots in each slice.

Canonical pathway enrichment (**Fig. 1E**) and correlation analysis (**Supplemental Fig. S1D)** revealed similarity between topics, with topics related to fibrotic tissue (topics 1, 13-14), regenerative or normal alveoli (4-6) or immune cells extravasation, immunoglobin and antigen presentation signaling (8 and 12). We then compared the average proportion of each topic in all spots of each lung section, indicating at day 14 large areas of fibrosis, enriched in fibroblast and macrophage markers (topics 1, 13-14) in lungs from both young and aged mice (**Fig. 1F**). At D28, lungs from young mice and, to a lesser extent, from aged mice, showed a reduced representation of topics related to fibrotic tissue compared to D14 while the areas corresponding to regenerative or normal alveoli (4-6) showed an opposite trend. Lung sections of old mice were also specifically enriched for the topic 8 (antigen presentation signaling), suggesting that infiltration of immune cells such as plasma cells, known to form aggregates in both IPF and the bleomycin model^28, 29^, is exacerbated by aging during fibrosis resolution. Overall, using various approaches, our data indicated that fibrosis resolution is delayed in lungs from aged mice.

### Time-resolved scRNA-seq captures cellular heterogeneity of bleomycin-injured lungs from young and aged mice

We reproduced the same experiment and performed scRNA-seq (10x Genomics Chromium platform) on whole lungs of young (7 weeks) and aged (18 months) mice (n=3) collected 14, 28 or 60 days after bleomycin challenge or PBS (**Fig. 2A**). The samples of both conditions from each time point were multiplexed using cell hashing^30^ to mitigate batch effects. The count matrices from the 36 sequenced samples were integrated to obtain a single dataset of 44,541 cells (**Supplemental Fig. S2A-B**). Based on their transcriptomic profile, cells were clustered and manually annotated based on the expression of specific gene markers (**Fig. 2B, Supplemental Fig. S2C and Supplementary table S2**). We identified 41 populations recapitulating the cellular heterogeneity of the mouse lung, of which 29 were immune cells from lymphoid or myeloid lineage, such as alveolar macrophages (AMs). We also detected mesenchymal cells, including fibroblasts, pericytes, mesothelial cells as well as epithelial populations such as AT1, AT2 and bronchial multiciliated cells. We finally identified 6 EC populations: proliferating EC (Prolif. EC), gCap, aCap, pulmonary-venous EC (PV EC), arterial EC and lymphatic EC. As expected, we observed a shift in several populations following bleomycin injury (**Fig. 2C and Supplementary table S3**). This was particularly the case for macrophages, as previously described ^31–33^, with a rapid influx of recruited monocyte-derived macrophages (Mo-AMs) AM3 populations along with a drop of the tissue-resident (TR-AMs) AM1 (**Supplemental Fig. S2D**). We evaluated the impact of age on the frequency of each AMs subpopulation relatively to the total number of cells analyzed in each mouse treated with PBS or BLM (**Supplemental Fig. S2E-F**). Overall, bleomycin treatment induced a very similar dynamics of macrophage subsets in young and old animals from the peak of fibrosis to complete resolution indicating that none of these specific subpopulations are associated with the delayed resolution in aged animals. Remarkably, bleomycin injury also induced major alterations in pulmonary EC types, especially gCap, PV EC and aCap (**Fig. 2C**), suggesting underlying cellular heterogeneity in bleomycin-treated mice.

**Figure 2.**
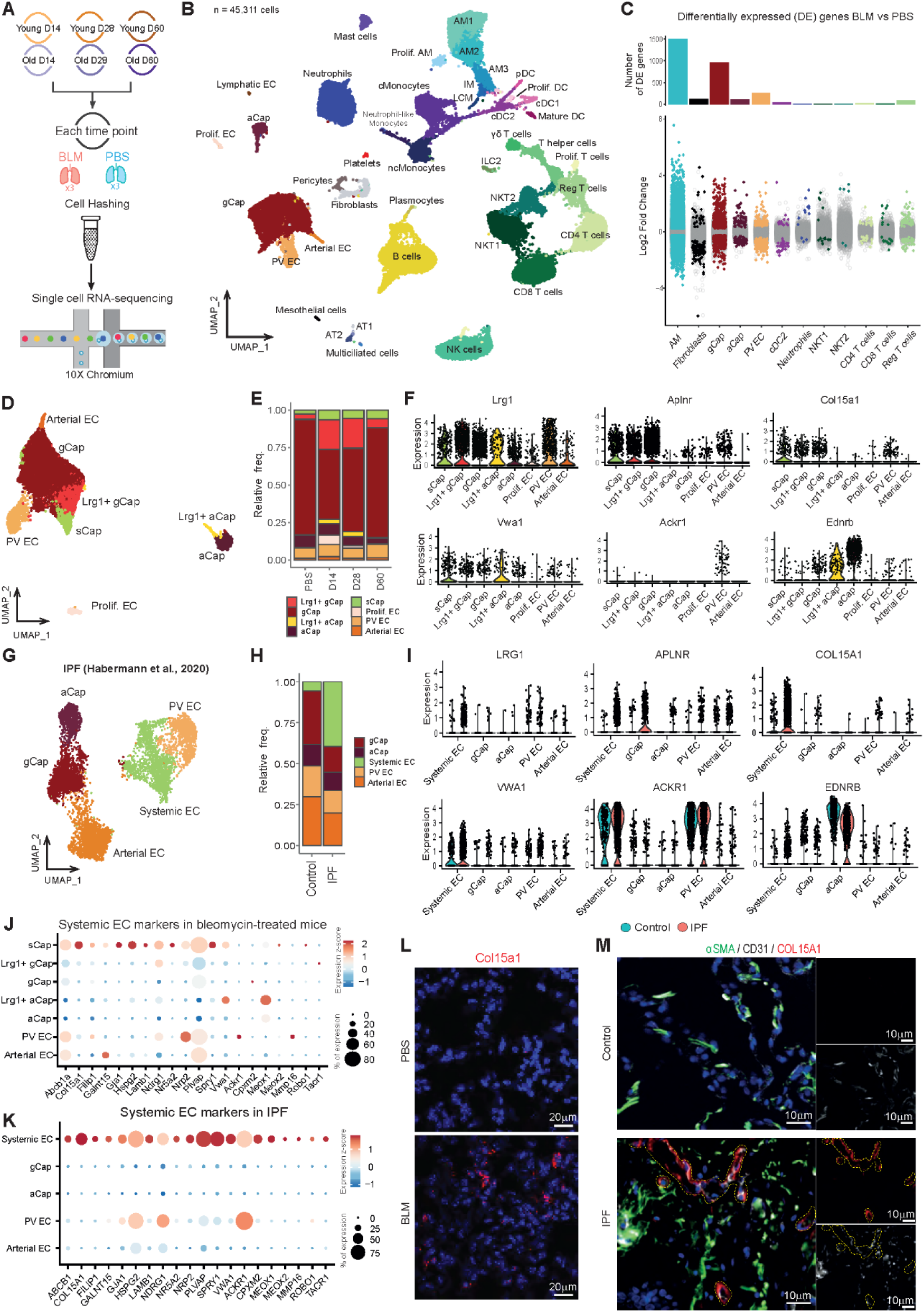
Time-resolved scRNA-seq captures pulmonary cell populations heterogeneity in bleomycin-injured lungs from young and old mice. **(A)** scRNA-seq experimental design (n=3, for each experimental condition at each time point). **(B)** UMAP of the integrated dataset of the 45311 sequenced cells. Cells are colored according to the annotated clusters. Prolif. AM = proliferating macrophages; AM = alveolar macrophages; IM = interstitial macrophages; LCM = large-cavity macrophages; Prolif. DC = proliferating dendritic cells; cDC = conventional dendritic cell; Mature DC = mature dendritic cells; pDC = plasmacytoid dendritic cells; cMonocytes/ncMonocytes = conventional/non-conventional monocytes; Prolif. T cells = proliferating T cells; Reg T cells = regulatory T cells; ILC2 = type 2 innate lymphoid cells; NKT = natural killer T cells; NK cells = natural killer; AT1/2 = Alveolar Type 1/2 cells. **(C)** Differentially expressed genes (DEGs) between Bleomycin and PBS treated cells for indicated populations. The number of DEGs is limited to 1500 for the bar plot. **(D)** UMAP of pulmonary EC subpopulations. gCap = general capillaries cells, aCap = aerocytes capillary endothelial cells, sCap = systemic capillary endothelial cells, PV EC = pulmonary-venous endothelial cells. **(E)** Relative proportions of PCEC subpopulations across time points. **(F)** Normalized expression of *Col15a1, Lrg1, Aplnr, Vwa1, Ackr1* and *Ednrb*. **(G)** UMAP of EC subpopulations from Habermann et al. dataset. **(H)** Relative proportion of EC subpopulations from Habermann et al. dataset. **(I)** Normalized expression of *LRG1, COL15A1, VWA1, ACKR1, APLNR* and *EDNRB* in EC subpopulations from control (blue) and IPF (light red) samples from Habermann et al. dataset. **(J-K)** Systemic EC markers comparison between mice **(J)** and human **(K)** fibrosis. (L) *In situ* hybridization of *Col15a1* mRNA in BLM or PBS conditions. **(M)** Expression of COL15A1 in lung sections from normal or IPF tissues by Immunofluorescence. Representative immunofluorescent images of COL15A1 (in red) with the pan EC marker CD31 (white) and αSMA (green). Nuclei are counterstained with DAPI. (n=5).

### Lung injury shifts PCEC towards Lrg1^pos^ subpopulations

The subclustering of PCEC and macrovascular subsets revealed the emergence of four new subpopulations in bleomycin-injured lungs from both young and aged mice (**Fig. 2D**). Among these subclusters, two new subpopulations of gCap and aCap cells emerged in fibrotic lungs with recognizable signatures particularly characterized by the overexpression of Leucine-rich α-2 glycoprotein 1 (*Lrg1*) and mostly found at day 14 and day 28 following bleomycin treatment (**Fig. 2E-F, Supplemental Fig. S3A and Table 1**). The third population matched a cluster of proliferating PCEC (Prolif. EC), which was only found at day 14 and to a lesser extent at day 28 following bleomycin treatment (**Fig. 2E and Supplementary table S2**). A fourth population expressed the “SV EC” marker Col15a1, an ectopic population, recently found widespread in fibrotic areas of IPF patients^13^. In order to establish a link between the endothelial remodeling induced after lung injury in the mouse model and human IPF, we re-analyzed the pulmonary endothelial subset from the Habermann dataset ^34^ (IPF Cell atlas), confirming the increase of the *COL15A1*-positive systemic EC (previously annotated as “SV EC”) proportion at the expense of others cell types in lungs from transplanted patients with IPF compared to healthy donors (**Fig. 2G-I).** The *Col15a1*^pos^ murine population shared, to some extent, the expression of others markers found in human systemic EC, such as *Hspg2, Nrp2, Gja1, Vwa1, Lamb1, Spry1, Filip1, Ndrg1, Nr5a2, Galnt15* and *Meox1* (**Fig. 2J-K and Supplementary table S4**). However, this population, described as being of macrovascular origin, doesn’t express macrovascular markers such as *Vwf* or *Vcam1* and also lacks expression of venous markers (Ackr1). Instead, the Col15a1^pos^ murine population shares several gCap markers such as *Aplnr* or *Sox11* (**Fig. 2F** and **Table 1**), confirming in mouse the identification of a systemic capillary ECs (sCap) recently characterized in lungs from lethal COVID-19 and IPF patients^15^. *In situ* hybridization confirmed the induction of *Col15a1* in BLM-treated mice (**Fig. 2L**), with a pattern similar as that observed in fibrotic areas of lung IPF sections (**Fig. 2M**). We hence considered these *Col15a1*^pos^ sCap as a distinct PCEC subpopulation for the rest of the study. On the other hand, we observed a low expression of *LRG1* in systemic EC from IPF samples (Habermann dataset^34^). We did not detect *LRG1* either in gCap or aCap from IPF samples, but decreased *EDNRB* was observed in aCap, as in aCap from bleomycin-treated mice **(Fig. 2F-I**).

**Table 1.**
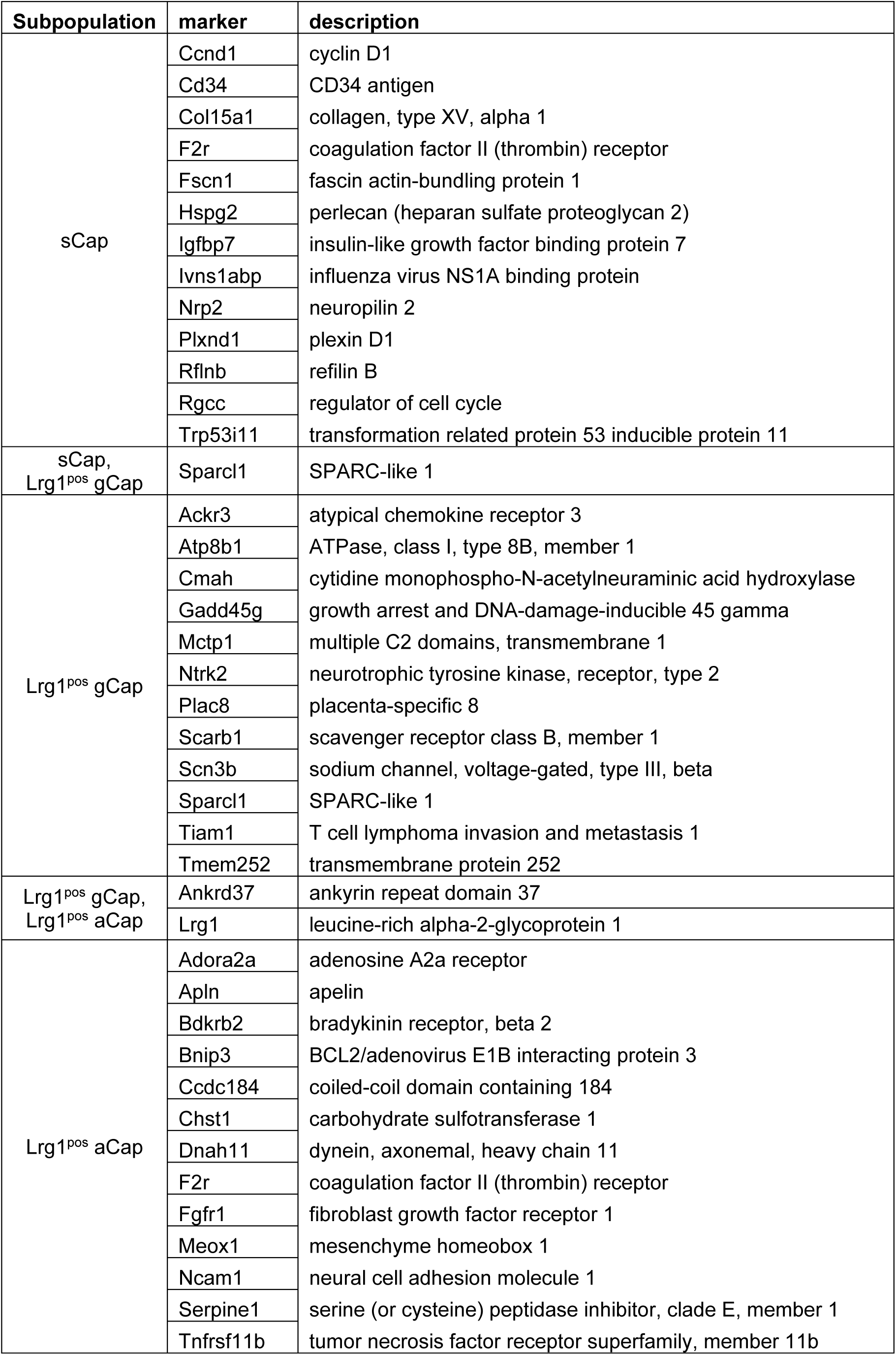
Main markers of Lrg1^pos^ PCEC subpopulations.

We then analyzed the profile of *Lrg1* expression in lung cell types in both basal condition or after BLM treatment in young and aged mice. In physiological condition, *Lrg1* expression is restricted to the macrovascular *Col15a1*-negative pulmonary-venous (PV EC), neutrophils, LCM, and to a lesser extent to AT2, gCap and Lymphatic EC **(Supplementary table S2** and **supplemental Fig. S3B).** At the peak of fibrosis, *Lrg1* expression was dramatically increased in all EC subpopulations as well as in alveolar macrophages subsets **(Supplemental Fig. S3B)**. We confirmed this increase of *Lrg1* in bleomycin-treated lungs by *in situ* hybridization **(Supplemental Fig. S3C)**. The kinetics of *Lrg1* expression in old versus young mice treated with BLM was similar in most subpopulations, apart for gCap, aCap and Arterial EC, for which *Lrg1* expression was time-shifted at day 28 **(Supplemental Fig. S3D).** Finally, overexpression of *Lrg1* in gCap and aCap was validated on an independent scRNA-seq dataset^35^ in which lungs of adult mice treated with bleomycin were collected at a shorter time interval **(Supplemental Fig. S3E).**

### Lrg1^pos^ PCEC signaling is associated with vascular remodeling and alveolar regeneration

*LRG1* is a gene coding for a pro-inflammatory glycoprotein, which can notably promote angiogenesis by switching TGF-β signaling from SMAD2/3 angiostatic signaling to SMAD1/5 angiogenic signaling^36^. To confirm this hypothesis, we generated a HMEC1 human endothelial cell line constitutively overexpressing LRG1. Control and LRG1 vector transduced cells were stimulated or not with TGF-β1 at 2 different concentrations (3 or 10 ng/ml). While in control cells, TGF-β1 mostly induced phosphorylation of SMAD2/3, LRG1 expression preferentially promoted SMAD1/5 phosphorylation at both low and high TGF-β1 concentrations (**Supplemental Fig. S4A-D**). This switch was also accompanied with a significant increase of *VEGF* transcript (**Supplemental Fig. S4E**), indicating that Lrg1^pos^PCEC signaling may participate in alveolar regeneration through induction of angiogenesis.

Ingenuity Pathway Analysis^TM^ performed on *Lrg1*^pos^ PCEC markers indicated an activation of mTOR, TGF-β, integrin-linked kinase and wound healing pathways in all 3 PCEC subpopulations (**Fig. 3A**). Enriched functions pointed to activation of angiogenesis or vasculogenesis, EC migration and proliferation in all 3 PCEC subpopulations, which is lacking or to a lesser extent in PV EC (**Fig. 3B**). We also noted elevated expression of hypoxia-regulated genes with *Lrg1*^pos^ aCap showing the strongest signature (**Fig. 3C**). Expression of markers associated with angiogenesis was shared by all three *Lrg1*^pos^ PCEC subpopulations, in particular genes coding for ECM-associated proteins, transcription factors/regulators, cytokines/growth factors and receptors. RNAscope^TM^ FISH confirmed the expression of HIF1α mRNA in *Lrg1*^pos^ cells in fibrotic areas, as well as increased expression of *Vegfa* and *Sox17* in gCaps at the peak of fibrosis (**Fig. 3E**). suggesting a central role for *Lrg1*^pos^ PCEC in endothelial regeneration processes via intense signaling within the alveolar niche (**Fig. 3D**).

**Figure 3.**
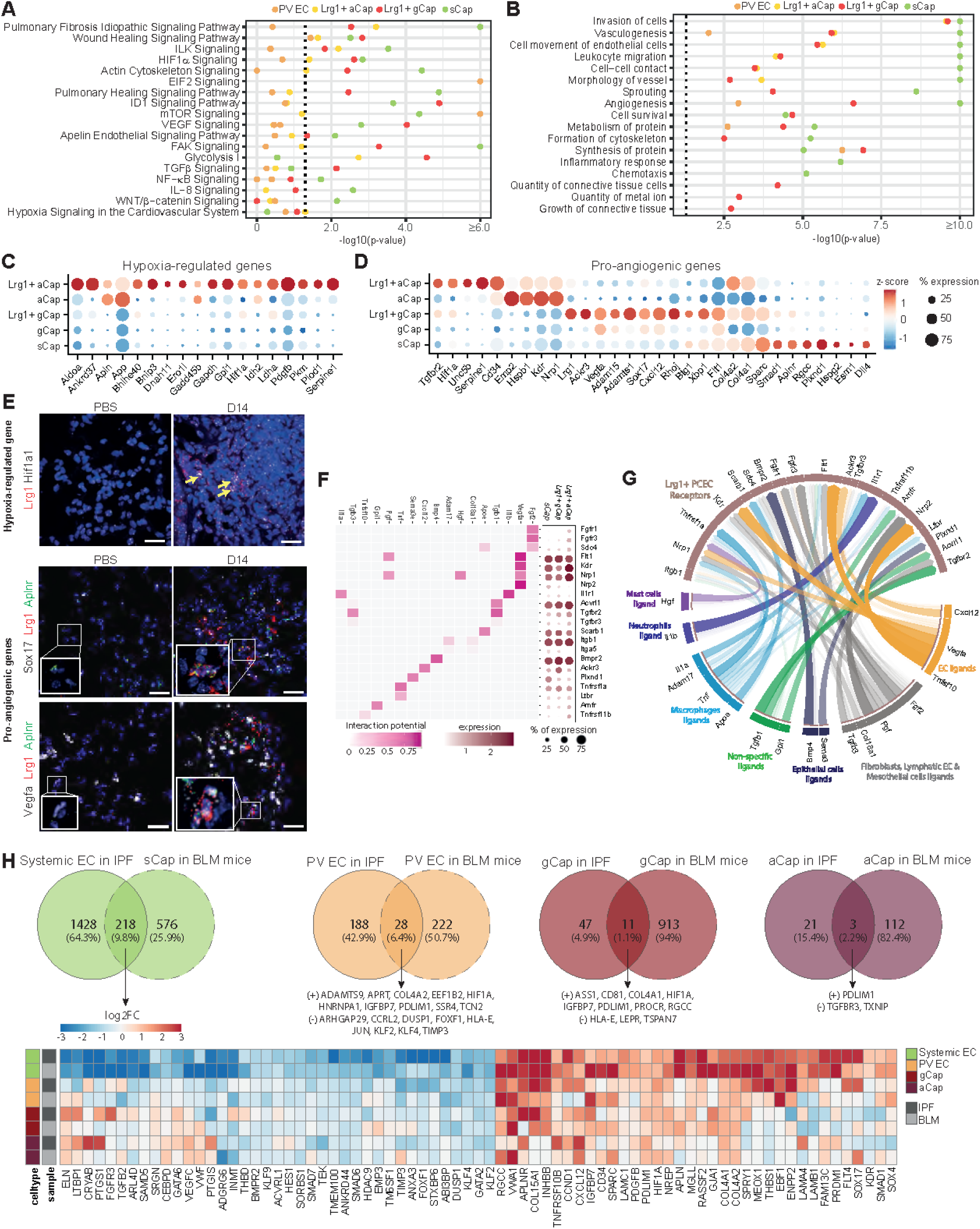
Lung injury shift pulmonary endothelial cells towards *Lrg1*^pos^ subpopulations associated with lung alveolar niche regeneration. **(A)** Selection of top activated canonical pathways in *Lrg1*^pos^ PCEC subpopulations. **(B)** Selection of top activated functions in *Lrg1*^pos^ PCEC subpopulations. **(C-D)** Level and percentage of expression of hypoxia-regulated (C) and pro-angiogenic (D) genes in *Lrg1*^pos^ PCEC subpopulations. **(E)** RNA FISH acquisition for *Hif1a1* (black, top), *Sox17* (black, middle), *Vegfa* (black, down), *Aplnr* (green) and *Lrg1* (red) mRNA in PBS or D14 in young mice. Scale bars =20µm. **(F)** Interaction potential between the prioritized ligands and their *bona fide* receptors (left); Level and percentage of expression of each receptor in *Lrg1*^pos^ PCEC subpopulations (right) **(G)** Circos plot of ligands-receptors interactions predicted by NicheNet analysis. Ligands are colored according to their specific expression. **(H)** Comparison of human and mouse pulmonary fibrosis signatures in Systemic EC, PV EC, gCap and aCap. Log2FC of similarly modulated genes in murine sCap and human systemic EC are displayed in the heatmap.

To follow up this hypothesis, we used NicheNet^37^ to infer ligand-receptor interactions potentially associated with the expression of genes related to the pro-angiogenic signature of Lrg1^pos^ PCEC. We selected the top ligands with the highest probability to induce these pro-angiogenic signatures in each Lrg1^pos^ PCEC subpopulation. Most of the selected ligands were commonly prioritized in the 3 subpopulations and were known pro-angiogenic factors, such as *Fgf2*, *Vegfa*, *Il1b*, *Tgfb1*, *Apoe*, *Bmp4*, *Adam17* or *Cxcl12* **(Supplemental Fig. S5A).** We next looked at the expression of the selected ligands as well as the downstream genes most likely to be regulated by these ligands in the different lung cell populations. Notably, *Vegfa* and *Cxcl12* ligands were among the top predicted ligands as well as the top predicted downstream-regulated genes, suggesting autocrine signaling within *Lrg1*^pos^ PCEC **(Supplemental Fig. S5A)**. We finally checked the expression of genes encoding *bona fide* receptors of the selected ligands in the 3 *Lrg1*^pos^ PCEC subpopulations (**Fig. 3F**). Overall, their transcripts were equivalently detected in the 3 *Lrg1*^pos^ EC subpopulations, with the exception of Fgf2 receptors Fgfr1 and Fgfr3 which are more specific to the *Lrg1*^pos^ aCap. We have summarized these interactions inferences in a circos plot, linking the selected ligands grouped by expression in specific cell populations to their top receptors expressed in *Lrg1*^pos^ PCEC subpopulations (**Fig. 3G**). We represented potential paracrine and autocrine signaling involving ligands expressed by mast cells (*Hgf*), neutrophils (*Il1b*), macrophages (*Il1a*, *Adam17*, *Tnf* and *Apoe*) as well as ligands expressed in multiple cell types such as *Tgfb1* and *Gpl1*. We also found potential interactions between ligands of neighboring cell types within the alveolar niche and *Lrg1*^pos^ PCEC receptors, such as the *Bmp4* / *Bmpr2* pair involving alveolar epithelial cells and the *Tgfb3* / *Tgfbr2* pair with mesenchymal cells. Based on the hypothesis that a receptor differentially expressed between physiological and fibrotic conditions likely indicates a modulation of the associated signaling pathway, we then compared receptor expression levels between gCap from bleomycin-versus PBS-treated mice. We observed that *Ackr3*, *Il1r1*, *Itgb1*, *Nrp2*, *Scarb1*, *Sdc4* and *Tgfbr2* were all upregulated while *Bmpr2* and *Tgfbr3* were downregulated in bleomycin-treated compared to control gCap **(Supplemental Fig. S5B)**. These observations appear consistent in the context of capillary endothelial remodeling. For example, the autocrine signaling pathway involving *Cxcl12* and *Ackr3* in PCEC has been shown to drive AEC expansion and neo-alveolarization by releasing Mmp14 following lung injury^38^. Our analysis indeed validated the upregulation of *Mmp14* in bleomycin-treated gCap at the single cell transcriptomic level **(Supplemental Fig. S5C)**. Finally, other genes driving key signal transduction proteins in EC, such as Eng, Id1 and several Smad family members, were also differentially expressed between both conditions in gCap **(Supplemental Fig. S5C)**. In particular, we found that *Smad6* was significantly downregulated in bleomycin-treated gCap. Coupled with the repression of *Bmpr2* and a similar trend for other BMP9 signaling pathway-associated genes (*Acvrl1*, *Eng*, and *Id1*), we hypothesized that this pathway responsible for EC quiescence may be inhibited in gCap following lung injury to promote angiogenesis^39^. Taken together, these results converge towards pro-angiogenic signaling in *Lrg1*^pos^ PCEC which, by releasing angiocrine factors, could thus contribute to the regeneration of the pulmonary alveolar niche.

### Bleomycin-induced endothelial remodeling echo pathological signatures found in human IPF

Using again the Habermann dataset^34^, we then systematically compared the mouse fibrotic signature induced by bleomycin with the IPF signature in systemic EC, PV EC, gCap, and aCap (**Fig. 3H and Supplementary table S6**). The most conserved pathological signature in both models was found for systemic EC, with a common modulation of 218 genes that includes most of the pro-angiogenic signature enriched in mouse *Lrg1*^pos^ PCEC (**Table 2**). Moreover, we observed a conserved upregulation of several signaling-associated genes, transcription factors and ECM-associated genes. With regard to conserved downregulated genes, we also observed a common inhibition of multiple transcription factors, including FOXF1, SMAD6 and GATA2 as documented recently^40^ as well as genes -associated with signaling, notably the BMP9 signaling genes (**Table 2**). These findings show that the pulmonary endothelial remodeling observed after bleomycin-induced lung injury in mice mimics at least in part the recruitment of systemic EC occurring in IPF, with a conserved signature associated with angiogenesis, ECM organization and cell adhesion.

**Table 2.** Commonly regulated genes in systemic EC from IPF samples and from bleomycin model.

| gene symbol (hsa) | description | function | expression_in_fibrosis |
| --- | --- | --- | --- |
| ACVRL1 | activin A receptor like type 1 | Angiogenesis | downregulated |
| APLN | apelin | Angiogenesis | upregulated |
| APLNR | apelin receptor | Angiogenesis | upregulated |
| BMPR2 | bone morphogenetic protein receptor type 2 | Angiogenesis | downregulated |
| FLT4 | fms related receptor tyrosine kinase 4 | Angiogenesis | upregulated |
| KDR | kinase insert domain receptor | Angiogenesis | upregulated |
| TEK | TEK receptor tyrosine kinase | Angiogenesis | downregulated |
| THBS1 | thrombospondin 1 | Angiogenesis | upregulated |
| TMEM100 | transmembrane protein 100 | Angiogenesis | downregulated |
| VEGFC | vascular endothelial growth factor C | Angiogenesis | downregulated |
| SRGN | serglycin | Apoptosis | downregulated |
| TNFRSF10B | TNF receptor superfamily member 10b | Apoptosis | upregulated |
| THBD | thrombomodulin | Blood coagulation | downregulated |
| CD34 | CD34 molecule | Cell adhesion | upregulated |
| IGFBP7 | insulin like growth factor binding protein 7 | Cell adhesion | upregulated |
| LAMA4 | laminin subunit alpha 4 | Cell adhesion | upregulated |
| LAMB1 | laminin subunit beta 1 | Cell adhesion | upregulated |
| LAMC1 | laminin subunit gamma 1 | Cell adhesion | upregulated |
| STXBP6 | syntaxin binding protein 6 | Cell adhesion | downregulated |
| VWF | von Willebrand factor | Cell adhesion,<br>Blood coagulation | downregulated |
| SORBS1 | sorbin and SH3 domain containing 1 | Cell adhesion, Cell signaling | downregulated |
| CCND1 | cyclin D1 | Cell cycle | upregulated |
| RASSF2 | Ras association domain family member 2 | Cell cycle | upregulated |
| RGCC | regulator of cell cycle | Cell cycle | upregulated |
| DUSP1 | dual specificity phosphatase 1 | Cell cycle, Stress response | downregulated |
| FGFR3 | fibroblast growth factor receptor 3 | Cell proliferation | downregulated |
| INHBB | inhibin subunit beta B | Cell signaling | upregulated |
| GJA1 | gap junction protein alpha 1 | Cell-cell communication | upregulated |
| ENPP2 | ectonucleotide<br>pyrophosphatase/phosphodiesterase 2 | Chemotaxis,<br>Lipid metabolism | upregulated |
| CXCL12 | C-X-C motif chemokine ligand 12 | Cytokine | upregulated |
| INMT | indolethylamine N-methyltransferase | Detoxification | downregulated |
| ABI3BP | ABI family member 3 binding protein | ECM organisation | downregulated |
| ADGRG6 | adhesion G protein-coupled receptor G6 | ECM organisation | downregulated |
| COL15A1 | collagen type XV alpha 1 chain | ECM organisation | upregulated |
| COL4A1 | collagen type IV alpha 1 chain | ECM organisation | upregulated |
| COL4A2 | collagen type IV alpha 2 chain | ECM organisation | upregulated |
| ELN | elastin | ECM organisation | downregulated |
| SPARC | secreted protein acidic and cysteine rich | ECM organisation | upregulated |
| VWA1 | von Willebrand factor A domain containing 1 | ECM organisation | upregulated |
| PDGFB | platelet derived growth factor subunit B | Growth factor | upregulated |
| TGFB2 | transforming growth factor beta 2 | Growth factor | downregulated |
| ARL4D | ADP ribosylation factor like GTPase 4D | Intracellular trafficking | downregulated |
| MGLL | monoglyceride lipase | Lipid metabolism | upregulated |
| PTGIS | prostaglandin I2 synthase | Lipid metabolism | downregulated |
| PTGS1 | prostaglandin-endoperoxide synthase 1 | Lipid metabolism | downregulated |
| TIMP3 | TIMP metalloproteinase inhibitor 3 | Metalloprotease inhibitor | downregulated |
| ANXA3 | annexin A3 | Phospholipase A2 inhibitor | downregulated |
| SPRY1 | sprouty RTK signaling antagonist 1 | Regulation of growth factor | upregulated |
| ANKRD44 | ankyrin repeat domain 44 | regulation of PP6 complex | downregulated |
| LTBP1 | latent TGFB binding protein 1 | Regulation of TGF-B | downregulated |
| NREP | neuronal regeneration related protein | Regulation of TGF-B | upregulated |
| CRYAB | crystallin alpha B | Stress response | downregulated |
| PDLIM1 | PDZ and LIM domain 1 | Stress response | upregulated |
| CEBPD | CCAAT enhancer binding protein delta | Transcription regulation | downregulated |
| EBF1 | EBF transcription factor 1 | Transcription regulation | upregulated |
| FOXF1 | forkhead box F1 | Transcription regulation | downregulated |
| GATA2 | GATA binding protein 2 | Transcription regulation | downregulated |
| GATA6 | GATA binding protein 6 | Transcription regulation | downregulated |
| HDAC9 | histone deacetylase 9 | Transcription regulation | downregulated |
| HES1 | hes family bHLH transcription factor 1 | Transcription regulation | downregulated |
| HIF1A | hypoxia inducible factor 1 subunit alpha | Transcription regulation | upregulated |
| KLF2 | KLF transcription factor 2 | Transcription regulation | downregulated |
| KLF4 | KLF transcription factor 4 | Transcription regulation | downregulated |
| KLF9 | KLF transcription factor 9 | Transcription regulation | downregulated |
| MEOX1 | mesenchyme homeobox 1 | Transcription regulation | upregulated |
| PRDM1 | PR/SET domain 1 | Transcription regulation | upregulated |
| SMAD1 | SMAD family member 1 | Transcription regulation | upregulated |
| SMAD6 | SMAD family member 6 | Transcription regulation | downregulated |
| SMAD7 | SMAD family member 7 | Transcription regulation | downregulated |
| SOX17 | SRY-box transcription factor 17 | Transcription regulation | upregulated |
| SOX4 | SRY-box transcription factor 4 | Transcription regulation | upregulated |
| EMP3 | epithelial membrane protein 3 | Transmembrane signaling | downregulated |
| FAM13C | family with sequence similarity 13 member C | Unknown | upregulated |
| SAMD5 | sterile alpha motif domain containing 5 | Unknown | downregulated |
| TM6SF1 | transmembrane 6 superfamily member 1 | Unknown | downregulated |

### *Lrg1*^pos^ PCEC dynamics are delayed in fibrotic aged mouse lungs

We then compared the dynamics of pulmonary EC after injury and during resolution between young and old mice (**Fig. 4A**), leveraging on their relative proportions found in each mice of the Chromium scRNA-seq dataset. The proportion of sCap, which were weakly detected in the control lungs, increased after bleomycin instillation and progressed during the resolution phase, suggesting that their recruitment is sustained over time independently of aging, even though the emergence of these cells appears to be more prominent in young mice at the peak of fibrosis. The presence of Col15a1^pos^ cells was confirmed by RNAscope^TM^ FISH in lungs from both BLM-challenged young and aged mice, from the peak of fibrosis up to D60 **(Fig. 4B**). We observed a similar profile between *Lrg1*^pos^ gCap and *Lrg1*^pos^ aCap, which sharply increased at d14 post bleomycin instillation and then diminishes upon resolution in young mice, while these increases were weaker and rather peaked at D28 in old animals. The simultaneous detection of Prolif. EC and conservation of the pool of physiological gCap and aCap supported that lung injury induces neovascularization of capillary EC. However, this regenerative response was stronger and shorter in young animals and also affected venous vessels.

**Figure 4.**
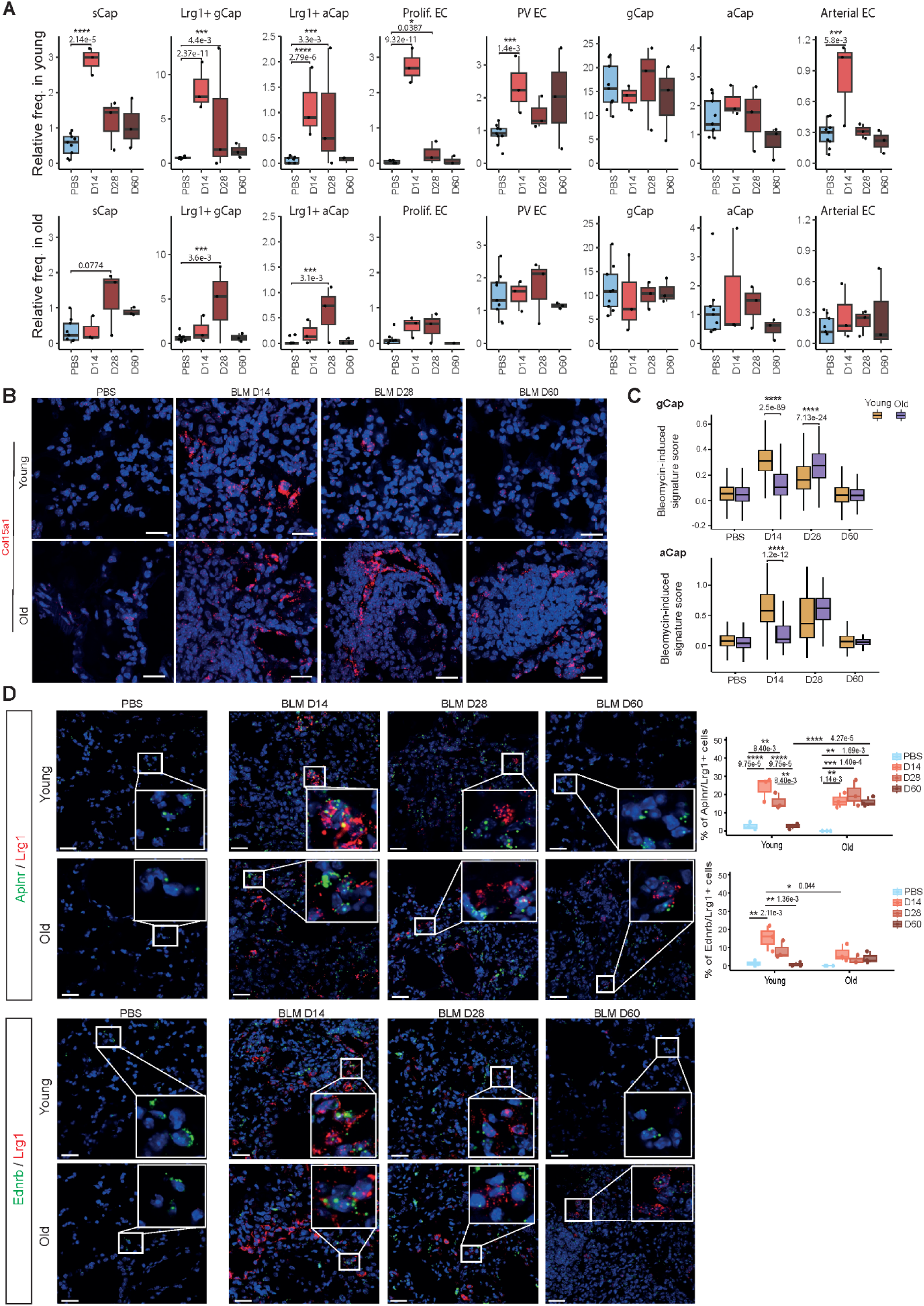
*Lrg1*^pos^ PCEC dynamics are delayed in old mice. **(A)** Pulmonary endothelial subpopulations relative frequencies in young and old mice across time points. **(B)** *In situ* hybridization of *Col15a1* mRNA at indicated time points in young and old mice. Scale bars = 20µm. **(C)** Bleomycin (BLM)-induced signature score (see material and methods section) in gCap (top) and aCap (bottom) in young and old mice across time points. **(D)** *In situ* hybridization of *Lrg1* mRNA with gCap marker *Aplnr* mRNA or aCap marker *Ednrb* mRNA at indicated time points in young and old mice. (n=3 independent mice, 3 counted microscopic fields / mouse, scale bars =20µm). Quantification was performed in three different fields for each mouse. * P<0.05, ** P<0.01, *** P<0.001, **** P<0.0001. P values were calculated by a Two-way ANOVA test followed by a multiple comparisons test with Holm-Sidak correction.

We confirmed these observations at the gene-level by calculating a score based on the expression of upregulated genes following bleomycin-induced lung injury, which we compared between cells grouped by age and time point (**Fig. 4C** and **Supplemental Fig. S6A-D**).

We also validated this delayed dynamic in an independent cohort of mice using *in situ* hybridization assay, showing that in young mice, *Lrg1^pos^* cells peaked at day 14 following bleomycin treatment and progressively decreased to reach their basal level at day 60, whereas a strong percentage was still maintained at day 28 and day 60 in old mice. Similarly, we observed the same evolution pattern with *Lrg1^pos^ Aplnr^pos^* cells and *Lrg1^pos^ Ednrb^pos^* cells, corresponding to *Lrg1^pos^* gCap and *Lrg1^pos^* aCap, respectively (**Fig. 4D**).

Finally, we returned to the spatial transcriptomic data to look for evidence of physiological and pathological PCEC signatures in spatial topics potentially associated with Lrg1 expression. Topic 6, highly represented in the young lung section at d28, was enriched in physiological gCap and aCap markers (*Sema3c, Hpgd, Thbd, Cldn5*, and *Car4*), as well as AT1 markers (*Ager, Hopx, Cldn18*), suggesting successful restoration of alveolar functions (**Fig. 1F, Supplemental Fig. S1B-C**). *Lrg1* was found in topics 1 and 4, where it was associated with fibroblast markers rather than endothelial markers. Interestingly, it was also enriched in topic 13, which displayed a clear inflammatory signature (**Fig. 1E**), notably carried by cytokines, complement and MHC components. This topic persisted at D28 in both lung sections from old mice (representing 6% and 10% of deconvoluted spots respectively), whereas it tended to be cleared (2% of deconvoluted spots) in the section from young mice at the same time point.

Taken together, our results showed that the dynamics of Lrg1^pos^ PCEC cells are delayed in lungs from aged mice, paralleling the retarded resolution of lung fibrosis observed in aged animals.

### Aged mouse lung gCap exhibit pro-inflammatory and injury-activated phenotype

We next performed differential expression analyses between old and young gCap in fibrotic or control conditions. In fibrotic conditions, we observed an enriched pro-inflammatory signature in old gCap, mainly characterized by the overexpression of genes encoding for MHC class II molecules, *Cd74*, *Cd52* and the interferon-induced protein Gbp4 (**Fig. 5A-B and Supplementary table S4**). The expression of *Cd74*, the receptor of Macrophage migration Inhibitory Factor (MIF), a critical upstream regulator of the immune response^41^ was validated by RNAScope^TM^ ISH, showing a stronger signal of *Cd74* in lungs from BLM-treated aged mice as well as partial co-localization of *Cd74* and *Lrg1* (**Fig. 5C**). Conversely, we found in young gCap an enrichment of transcription factors associated with vascular homeostasis and repair such as *Klf2, Klf10 and Peg3*^42–44^ (**Fig. 5B and Supplementary table S4**). In PBS-treated mice, comparison between old and young gCap revealed even more gene modulations (**Fig. 5D-E**), which were associated with multiple functions, including an increase of inflammatory and immune responses, but also ECM organization and IPF signaling pathway in aged mice (**Fig. 5F and Supplementary table S5**). Indeed, this aging signature contained 21 genes significantly deregulated in gCap following bleomycin challenge, including the top *Lrg1*^pos^ PCEC markers *Lrg1*, *Col15a1*, *Nrtk2*, *Ndrg1* and *Ankrd37* (**Fig. 5G**), indicating that old gCap in physiological condition already express a pro-inflammatory and injury-activated signature. We also observed that old gCap overexpressed *Vwf*, *Slc6a2*, *Prss23 and Plat* (**Fig. 5D-E**), which are normally restricted to macrovascular EC **(Supplemental Fig. S3A).** These cells also show a decrease in *Aplnr* expression compared to young gCap, in line with a predicted decrease in the Apelin signaling pathway in old gCap (**Fig. 5F**), strongly suggesting a potential change in their intrinsic functions.

**Figure 5.**
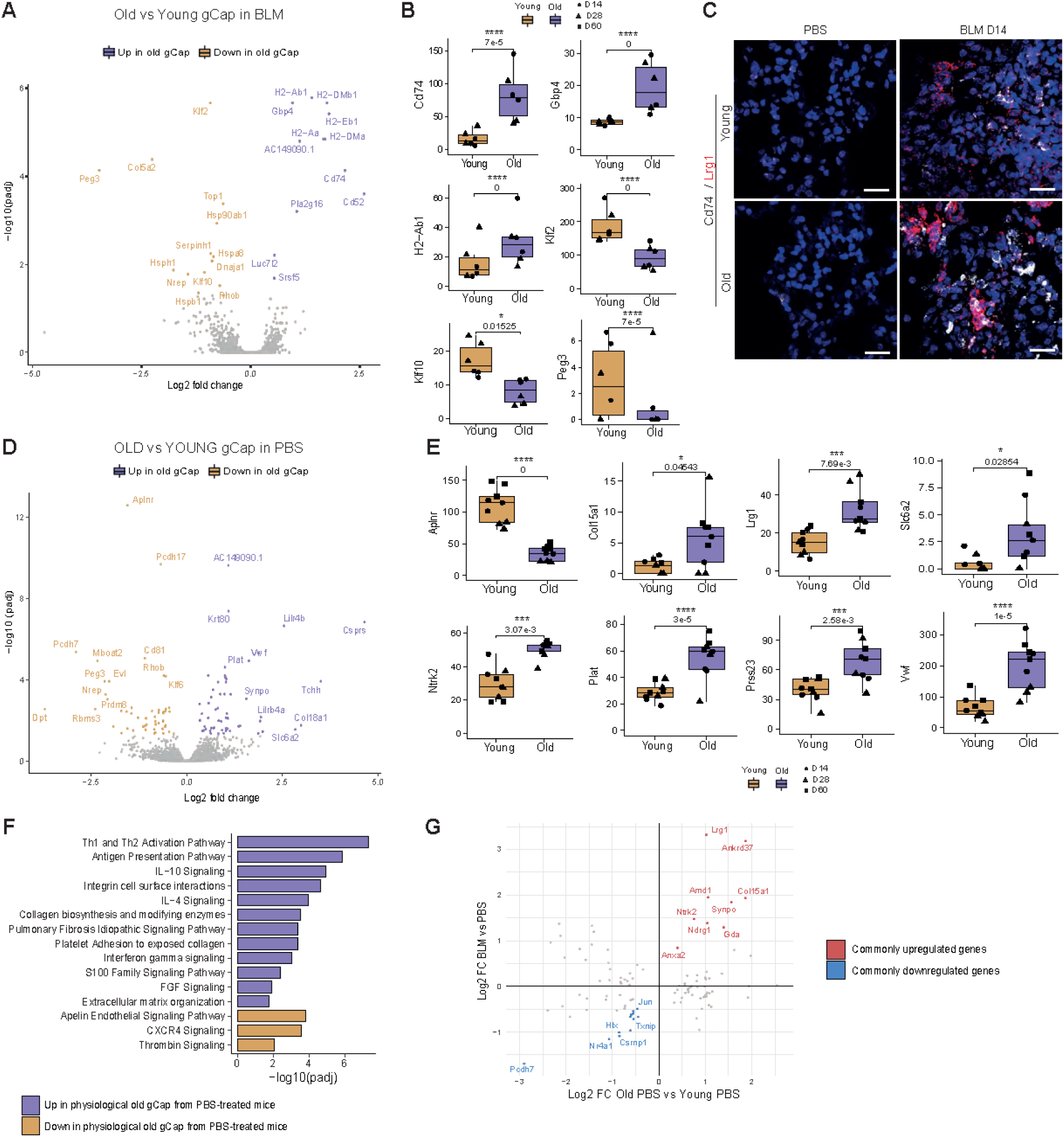
Aged mouse lung gCap exhibit pro-inflammatory and injury-activated phenotype. **(A)** Differentially expressed genes (DEGs) between old and young gCap in fibrotic condition. **(B)** Expression of several genes dysregulated in age-related in gCap under BLM conditions (*Cd74, Gbp4, H2-Ab1, Klf2, Klf10* and *Peg3*). Boxplot of relative expression, each point corresponds to a mice (● for D14, ▴ for D28 and ▪ for D60). **(C)** *In situ* hybridization of *Cd74* mRNA in BLM or Day 14 after bleomycin induction in young and old mice. Scale bars = 20µm. **(D)** DEGs between old and young gCap in physiological condition. **(E)** Expression of several genes dysregulated in age-related in gCap under PBS conditions (*Aplnr, Col15a1, Lrg1, Slc6a2, Ntrk2, Plat, Prss23* and *Vwf*). Boxplot of relative expression, each point corresponds to a mice (● for D14, ▴ for D28 and ▪ for D60). **(F)** Function enrichment analysis on DEGs between old and young physiological gCap. (**G**) Expression of several age-related dysregulated genes in gCap under BLM/PBS conditions.

### Trajectory inference indicates a transition between *Col15a1*^pos^ sCap, *Lrg1*^pos^ gCap and physiological gCap following alveolar injury

The parallel kinetic profile between the *Lrg1*^pos^ PCECs populations after BLM treatment as well as their transcriptomic proximity and delayed appearance in aged animals, correlating with a delayed resolution, suggest that a dynamic process links these different PCEC subpopulations or cell states during alveolar niche regeneration. To model a potential trajectory in an unbiased way, we used the generalized RNA velocity approach^45^, leveraging on the splicing dynamics estimated in young bleomycin and PBS-treated mice samples at day 14. Latent time ordering indicated that *Col15a1*^pos^ sCap could initiate a differentiation process, transiting through *Lrg1*^pos^ gCap before ending up to physiological gCap **(Fig. 6A**). Of note, the same trajectory was proposed in the same configuration in the old mice 28 days after instillation (**Fig. 6B**). This inferred trajectory was supported by genes displaying a dynamic expression profile according to latent time, either progressively decreasing (*Smad1, Sparc, Col4a1, Ndrg1, Igfbp7*), transiently upregulated (*Ackr3, Emp1, Cmah, Vwf, Smad7*), or progressively increasing (*Npr3*, *Hpgd*, *Clec1a*, *Bmp6*, *Itga1*) (**Fig. 6C-D and Supplemental Fig. S7A)**. These data suggest that in the bleomycin reversible fibrosis model, sCap are recruited to contribute to gCap replenishment and to the regeneration of the lung capillary endothelium integrity (**Supplemental Fig. S7B).**

**Figure 6.**
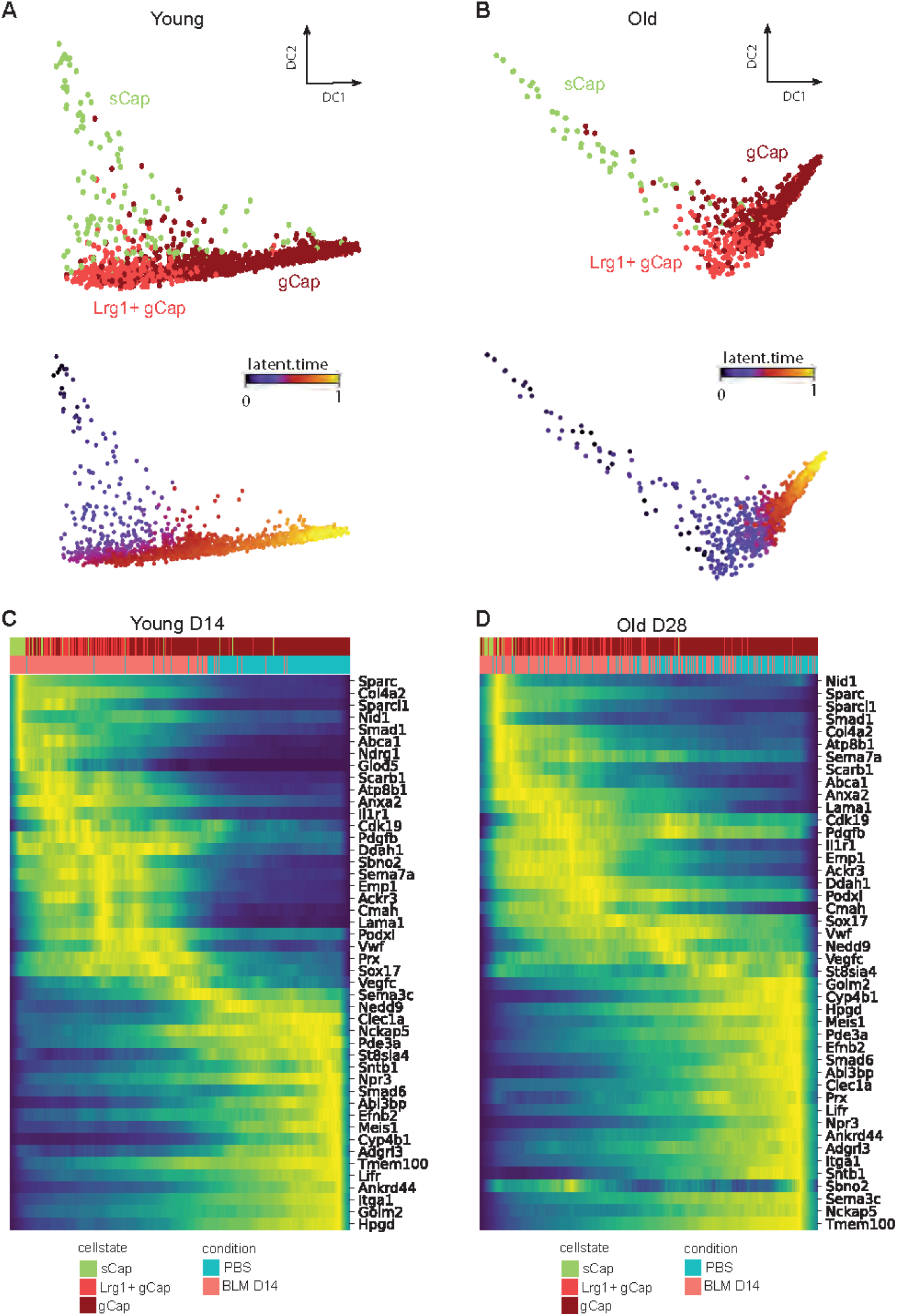
RNA velocity analysis reveals transcriptional dynamics of PCEC during alveolar regeneration. **(A-B)** Diffusion map computed on Young D14 **(A)** or old D28 **(B)** mouse spliced RNA data (3 Bleomycin and 3 PBS-treated mice), with cells colored according to PCEC subpopulation (up) and RNA velocity latent time (bottom). **(C-D)** Heatmap of the top contributing genes to the transcriptional dynamic, with cells ordered by latent time in young D14 **(C)** or old D28 **(D)** mice.

## DISCUSSION

In this work, we used scRNA-seq combined with spatial transcriptomics to follow the transcriptomic modifications of lung cell populations during fibrosis formation and resolution in a mouse model of lung fibrosis. We specifically focused on the transcriptomic alterations correlated with a delayed fibrosis resolution in aged mice and identified several aging-associated changes in some PCEC subpopulations, including aCap and gCap, shifting towards a pro-angiogenic phenotype expressing *Lrg1* and associated with alveolar niche regeneration. We also found that aging altered the transcriptome of gCap in control mice, with a signature suggestive of a pro-fibrotic and injury-activated state. Because PCEC are intimately connected to alveolar epithelial cells and fibroblasts in the alveolus, age-associated defects in the PCEC differentiation may likely participate in the dysfunction of other cells in the alveolar niche and impede alveolar regeneration.

Our study presents a whole picture of cell dynamic changes in lung from young and aged animals during fibrosis formation and resolution at 3 time points. Using several approaches, we confirmed previous works showing that aged mice exhibit impaired resolution after bleomycin injury^23–25^, further proposing using spatial transcriptomics that infiltration of immune cells such as plasma cells, known in IPF to form aggregates and bronchus-associated lymphoid tissue (BALT)^29^, contribute to this delayed resolution in aged mice. We also confirmed by scRNA-seq that many cell populations are affected by bleomycin, including AEC, fibroblasts, macrophages and PCEC^26, 32, 33, 46^, providing a useful dataset to analyze the transcriptomic dynamic of these populations and compared it to the kinetic of the resolution process measured with histology, collagen production and spatial transcriptomic.

One of the most affected cell type by the bleomycin challenge in our dataset corresponded to EC, with several main subpopulations that were only apparent in bleomycin-injured lungs. First, we identified a homologous Col15a1-positive systemic EC population, previously described in lungs from IPF patients^13^. We also found that this population, described originally as being of venous macrovascular origin, doesn’t express clear macrovascular and venous markers and likely corresponded to a systemic capillary ECs (sCap) in agreement with a recent study in COVID-19 patients^15^. Secondly, we found two new subpopulations of gCap and aCap cells in fibrotic lungs sharing a partial common gene signature characterized by the overexpression of *Lrg1*. Interestingly, among these distinct populations, only *Lrg1*^pos^ gCap and *Lrg1*^pos^ aCap cells displayed a different kinetic profile following bleomycin challenge in young and aged mice. While they both peaked at day 14 in young mice lungs, their dynamic was strongly shifted in the lungs of aged mice, peaking at day 28. Overall, our findings demonstrate that bleomycin injury induces PCEC remodeling marked by the upregulation of *Lrg1* and the emergence of distinct capillary activated cell states that are delayed in aged mice.

*Lrg1* codes for a secreted member of the family of leucine-rich repeat proteins, that often served as pattern recognition motifs for the innate immune system^47^. It is a multifunctional signaling molecule initially associated with pathological angiogenesis^36^ that can notably modulate the TGF-β pathway in a highly context-dependent manner. The physiological function of *Lrg1* remains poorly understood, with knockout mice showing no obvious phenotypic defects. Several studies indicate that it promotes physiological wound healing and maintain tissue homeostasis^48, 49^, although abnormal expression levels has been found to disturb effective would healing and contribute to fibrosis in several tissues including lung^50^. These data suggest that *Lrg1* expression must be finely regulated during the repair process and that abnormal levels may disrupt effective wound healing and contribute to fibrogenesis. One of the most compelling examples of LRG1 function in promoting diseased vessels is commonly described as the “TGF-β angiogenic switch”, which switches TGF-β signaling to a proliferative pathway in a variety of pathological settings, including age-related macular degeneration and cancer^36^. While binding of TGF-β to endothelial TGF-β type II receptor (TGFβRII) normally initiates signaling through the tyrosine kinase receptor ALK5 coupled to SMAD2/3, preserving cell quiescence, high levels of LRG1 can redirect TGF-β to form a transduction complex with ALK1 and the accessory receptor endoglin (ENG), activating the pro-angiogenic SMAD1/5/8 pathway and promoting EC proliferation, migration and tubulogenesis as well as the expression of several pro-angiogenic factors such as VEGFA. Our in vitro experiments confirmed this molecular mechanism, which likely explains our data showing inhibition of endothelial quiescence in gCap *Lrg1*^pos^ as well as strong induction of markers associated with EC proliferation and migration after lung injury to promote angiogenesis and sprouting. During the preparation of this manuscript, Zhao et al. indeed demonstrated that *Lrg1* was able to counterbalance the canonical angiostatic TGFβRII signaling to promote lung vascular repair following viral injury^51^. While this *Lrg1*-associated pro-angiogenic activity appears to be necessary to promote alveolar vascular repair, it is likely that delayed expression of *Lrg1* in aged mice could slow down the formation of a new mature capillary network thus causing a delay in the regeneration of the alveolar niche.

The *Lrg1*-associated gene signature found in our study also contains *Sox17*, a member of the Sry-related high mobility group domain family F (Sox F) transcription factors, a key developmental regulator of EC lineage. Previous work indicated that induction of *Sox17* expression through the activation of HIF-1α in EC plays an essential role in regenerating EC following endotoxin-induced EC injury, notably by transcribing Cyclin E1 and stimulating proliferation of EC^52^. Of note, both *SOX17* and *HIF1A* were also detected in PCEC in lungs from IPF patients, indicative of a pro-angiogenic activity. Overexpression of these two genes were validated in fibrotic areas of lung mice, with co-localization with *Aplnr* or *Lrg1*, respectively. The activation of a hypoxic response was also particularly visible in *Lrg1*^pos^ aCap and *Lrg1*^pos^ gCap with several markers associated with glycolysis and hypoxia, in agreement with studies indicating that transient activation of hypoxia-mediated signaling pathways contributed to angiogenesis and vascular repair^53^. In addition to hypoxia, our analysis revealed transcriptomic modulations of mTOR, TGF-β, IL-8 and integrin-linked kinase signaling and related notably to vasculogenesis, sprouting, cell-cell contact, cell migration and inflammation. The identification of multiple cytokines/growth factors and receptors in activated PCEC suggested extensive heterotypic interactions within the alveolar niche. Our ligand-receptor interaction predictions propose a complex network of interactions involving ligands expressed by activated PCEC and neighboring cell types within the alveolar niche, including AT1, AT2, mesenchymal cells as well as mast cells, neutrophils and macrophages that likely influence alveolar repair. Remarkably, some of these predictions are in agreement with previous studies highlighting in particular the importance of the signaling pathway involving *Cxcl12* (also known as SDF-1) and *Ackr3* (*Cxrc7*) in PCEC, leading to neo-alveolarization^38^. Further work is needed to analyze the functional importance of this predicted network and to characterize their central regulators with the goal to potentially target specific signaling pathways within the vascular niche for regenerative therapies.

We also found additional aging-related transcriptomic differences in PCEC subpopulations in physiological or pathological conditions. We focused on gCap population that functions as progenitor cell for aCap in capillary homeostasis and repair^9^. We found that in lungs from aged bleomycin-challenged mice, gCap expressed an enriched pro-inflammatory and immune signature, as well as decreased expression levels of several transcription factors associated with vascular homeostasis, repair and the anti-inflammatory response (*Klf2, Peg3, Klf10)*^42–44^, indicating that the bleomycin response was exacerbated in gCap from aged mice. In addition, we also found an aging-related signature in gCap from healthy control lungs. Part of this signature (21 genes), corresponding mostly to genes associated with angiogenesis, migration and sprouting, was correlated with the bleomycin-induced signature, suggesting a pre-inflammatory and injury-activated cell state in old gCap.

Our study has limitations, in particular because of some technical biases during the single-cell and spatial transcriptomic workflow. First, our dissociation protocol, combined with the cell Hashing protocol^30^, requiring additional cell labeling and washes has led to a depletion of fibroblasts and epithelial cells, which did not allow us to perform a complete cell state analysis for these cell types. Secondly, while the Visium spatial technology skips the dissociation step, its resolution did not allow to perform a single-cell characterization of the multiple cell lineages composing the alveolar niche, despite the use of a deconvolution tool. Moreover, although this approach was intended to highlight some spatial information explaining the resolution delay in aged mice, it will need to be supplemented by a larger number of replicates to improve these observations. Finally, the bleomycin mouse model may not accurately recapitulate the vascular alterations observed in IPF patients. The comparison of our data with a public dataset from the IPF cell atlas^34^ indicated that *LRG1* transcript was expressed at a low level in *COL15A1*^pos^ systemic EC, questioning the relevance of this mouse model. Nevertheless, several conclusions could be drawn from this analysis, including a large conserved pathological signature of 218 genes between the *Col15a1*^pos^ sCap from fibrotic mouse and IPF lungs, confirming their close functional state and a common pro-angiogenic signature between PCECs from mouse fibrotic lungs and human IPF systemic EC, suggesting that the pulmonary endothelial remodeling observed after bleomycin-induced lung injury in mice mimics the recruitment of systemic EC occurring in IPF. In contrast, the aCap *Lrg1*^pos^ and gCap *Lrg1*^pos^ populations were not clearly detected in IPF samples and only a small fraction of genes appeared to be commonly modulated in aCap/gCap populations from fibrotic lungs in humans and mice. The characterization of EC heterogeneity in IPF is still in its infancy. In particular, it is not known whether the ectopic presence of *COL15A1*^pos^ systemic EC in the IPF-associated distal parenchyma, play a role in the pathogenesis of IPF. In order to gain information on their potential function, we used our scRNA-seq dataset on young PCEC at day 14 to investigate the potential relationship between sCap and gCap using the RNA velocity approach^45^. Our results indicate that following lung injury, sCap are recruited and may function as progenitor cells that can initiate a differentiation process, transiting through Lrg1^pos^ gCap before reaching the physiological gCap cell state. The same differentiation scheme was also found in the old mice but with a delayed kinetic (28 days). Overall, these data suggest that in a reversible fibrosis model (mouse), sCap are recruited and can act as progenitor cells to reconstitute the gCap pool and regenerate the integrity of the alveolar endothelial capillaries while aging appears to delay this resolution process. It is therefore tempting to hypothesize that in the context of the human pathology, systemic EC might be deficient in this differentiation process and accumulate in the injured tissue. Interestingly, a recent article, using an *ex vivo* human precision-cut lung slice (hPCLS) model of lung fibrogenesis, also described a transcriptomic proximity between alveolar PCEC and a systemic venous EC state^54^. In this case, the authors presented a differentiation trajectory in the opposite direction, proposing that PCEC in injured alveoli give rise to a *VWA1*^pos^/*PLVAP*^pos^ ectopic EC state with transcriptional similarities to the systemic vasculature. Additional analyses are thus needed to better characterize this ectopic EC state versus the *COL15A1*^pos^ systemic EC and distinguish between these two hypotheses. During the revision of this manuscript, a study by Raslan et al. also investigated the bleomycin-induced lung injury response in young and aged mice at single-cell resolution^55^. Interestingly, they also isolated a sub-population of gCap EC marked by the expression of *Ntrk2/TrkB* (Tropomyosin Receptor Kinase B) that appeared in bleomycin-injured lungs and associated with aberrant YAP/TAZ signaling, mitochondrial dysfunction, and hypoxia. These cells exhibited a similar gene expression signature as those we present here, including the upregulation of *Lrg1*. However, their conclusions, stating that this population is dysfunctional and accumulates with aging, differ from ours. One explanation for this discrepancy could be that they studied only a single time point (day 30), without considering the dynamic of appearance and disappearance of this population at earlier and later times. Further functional investigations aimed at depleting these *Lrg1*^pos^ sub-populations *in vivo* will be required in order to clarify these two contrasting hypotheses.

Altogether, our findings shed new light into the molecular mechanisms associated with alveolar endothelial capillaries resolution in a mouse model of reversible lung fibrosis and how aging influences specific PCEC to delay this resolution process. In particular, they raise new hypotheses regarding the functional importance of these specific PCEC populations in lung repair, which may lead to new therapeutic strategies for the treatment of IPF.

## METHODS

### Reagents and antibodies

Bleomycin was obtained from Sigma Aldrich (B1141000). The following antibodies were used in immunofluorescence experiments: Mouse Anti-Human CD31 (MA5-13188 Thermofisher), Rabbit Anti-Human COL15A1 (53667 Invitrogen), Purified Mouse Anti-αSMA–FITC (F3777 Sigma-Aldrich), donkey DyLight 594 Anti-Rabbit IgG (A32754 Life Technologies), donkey DyLight 647 Anti-Mouse IgG (A31571 Life Technologies). The following antibodies were used in western blot experiments: Rabbit anti-Human pSmad2 (18338, Cell signaling), Rabbit anti-Human Smad2 (5339, Cell signaling), Rabbit anti-Human pSmad1/5 (9516, Cell signaling),Rabbit anti-Human Smad5 (9517, Cell signaling), Rabbit anti-Human LRG1 (PA5-96832, Thermofisher), Rabbit anti-Human HSP90 (4877, Cell signaling) and Goat anti-rabbit HRP (P0448, Dako).

### Experimental design and animal treatment

All animal care and experimental protocols were conducted according to European, national and institutional regulations (Protocol numbers: 00236.03 and APAFIS#31298, IPMC approval E0615252; Protocol Number APAFIS#12540, University of Lille; approval 00236.03 CNRS). Personnel from the laboratory performed all experimental protocols under strict guidelines to ensure careful and consistent handling of the mice. Seven-week (“young”) and 18-month (“old”)-old C57BL/6 male mice (Charles River) were divided randomly into two groups: (A) saline-only (PBS, n=3), or (B) bleomycin (BLM, n=3). To induce fibrotic changes, 50-µl bleomycin (1 U/kg) or PBS was aerosolized in mouse lungs using a MicroSprayer Aerosolizer (Penn-Century, Inc.) as previously described ^56^. Mice were sacrificed at designated time points (days 14, 28 and 60) after instillation.

### IPF lung tissue sections

Lung tissue sections were obtained from the UGMLC Giessen Biobank affiliated to the European IPF Registry as well as from the Nice Hospital-Integrated Biobank (transfer authorization from the French ministry of research n° AC-2021-4655).

### Histopathology

Lungs were fixed overnight with neutral buffered formalin and then embedded in paraffin. 5-micrometer-thick sections were mounted and stained with hematoxylin and eosin (HE) as well as Sirius Red to assess the degree of fibrosis. HE sections were scanned using Axio Scan Z1 scanner (Zeiss) at x10 magnification. Data are processed using the Zen blue imaging software (Zeiss). Hematoxylin and eosin zoomed in pictures were performed using Axiophot 2 light microscope.

### Hydroxyproline quantification of mouse lungs

Lung hydroxyproline content was assayed as previously described ^56^. Briefly, lungs were disrupted in liquid nitrogen and 10 mg were homogenized in 100 µl of water. Samples were hydrolysed for 3 hours at 120°C in a temperature-controlled heating block after adding 100 µl of HCl 12M and then dried. Hydroxyproline quantification was finally performed in microplate as described by the manufacturer (Sigma).

### Visium^TM^ spatial transcriptomics protocol

#### Tissue preparation

Tissue preparation and slides processing were performed according to the Visium Tissue Preparation Guide (CG000238 Rev A; CG000239 Rev A; CG000240 Rev A, 10× Genomics). Briefly, lungs of bleomycin-challenged mice (Young.D14/D28; Old.D14/D28, n=2 for a total of 8 samples) were inflated with diluted Optimal Cutting Temperature (OCT), gently removed from mice and then embedded in OCT on dry ice and stored at −80°C. OCT blocks were cryosectioned at 10µm thickness on capture areas of Visium 10 Genomics slide. Before performing the complete protocol, Visium Spatial Tissue Optimization was performed according to manufacturer’s procedure and using the Axioscan 7 scanner (Zeiss). Thirty minutes was selected as optimal permeabilization time. The experimental samples were fixed, stained with H&E and imaged using Axioscan 7 scanner at 5x magnification.

#### Generation of spatial transcriptomic libraries

Library preparation was performed according to the Visium Spatial Gene Expression User Guide. Libraries were loaded and sequenced on a NextSeq2000 System (Illumina) as paired-end-dual-indexed, at a sequencing depth of approximately 25 M read-pairs per capture area.

#### Analysis of spatial transcriptomics data

The 8 space ranger pipeline outputs containing both the spot-level gene counts and the histological image of the tissue slice were read as seurat objects. After a quick overview of each one, 2 of them (Young.D14_#2 and Young.D28_#2) appeared unusable because of a very low level of detected molecules per spot (<800 detected genes). The remaining 6 samples (Young.D14, Old.D14.a, Old.D14.b, Young.D28, Old.D28.a, Old.D28.b) were jointly processed using the SpatialExperiment^57^ (v1.10.0) and STDeconvolve^27^ (v1.3.1) R packages. Firstly, the genes x spots count matrices of the 6 samples were stripped of mitochondrial, ribosomal, unannotated non-coding and hemoglobin subunit coding genes. The preprocess() function was run a first time on each sample to identify overdispersed genes, using the following arguments: nTopGenes = 5, removeAbove = 0.95, min.reads = 10, min.lib.size = 1, min.detected = 1, ODgenes = TRUE, od.genes.alpha = 0.05, gam.k = 5, nTopOD = 1000. We took the union of the 6 lists of overdispersed genes, representing 2482 unique genes. The preprocess() function was run a second time on the concatenated count matrix of those 2482 genes in all spots, specifying to discard spots containing less than 200 UMIs, and genes detected in less than 10 spots, resulting in a corpus of 2479 genes × 10813 spots count matrix and the associated spatial coordinates. The fitLDA() function was run on the count matrix in order to help selecting an optimal number K of predicted topics. Following the recommendations of the authors, we chose a value of K=14, i.e. the value that produces the smallest perplexity without exceeding 5 topics with an average proportion of less than 5%. The topics proportions across spots (theta) and topics gene probabilities (beta) matrices from the optimal fitted LDA model were obtained with the getBetaTheta() function with a betaScale factor = 1000 to scale the predicted topics gene expression profiles. To annotate each of the 14 topics, the log2FC of all expressed genes were calculated by dividing the expression in the topic versus the average expression in the rest of the topics. Genes with a log2FC > 0.4 were used to perform a canonical pathway enrichment in each topic using Ingenuity Pathway Analysis (Ingenuity® Systems, www.ingenuity.com).

### Chromium^TM^ single-cell transcriptomics protocol

#### Lung dissociation and cell preparation

Lung single cell suspensions were generated as previously described^58^. Briefly, after euthanasia, lung tissue was perfused with sterile saline through the heart and were inflated through the trachea with dissociation cocktail containing dispase (50 caseinolytic U/ml), collagenase A (2 mg/ml), elastase (1 mg/ml), and DNase (30 μg/ml). Lungs were immediately removed, minced to small pieces ∼1mm3), and transferred for mild enzymatic digestion for 20–30 min at 37 °C in 4 ml of the dissociation cocktail. Enzymatic activity was inhibited by adding 5 mL of PBS supplemented with 10% fetal calf serum (FCS). Single cells suspensions were passed through a 40-micron mesh, and then harvested by centrifugation at 300 g for 5 min (4°C). Reb blood cell lysis was performed using RBC lysis buffer (Thermo fisher) for 2 min at 4°C and stopped using PBS 10% FCS. After another centrifugation at 300g for 5 min (4°C) cells were counted and critically assessed for single cell separation and overall cell viability using the Countess 3 FL (Fisher Scientific). Samples were then stained for multiplexing using cell hashing [3], using the Cell Hashing Total-Seq-ATM protocol (Biolegend) following the protocol provided by the supplier, using 6 distinct Hash Tag Oligonucleotides-conjugated mAbs (3 PBS and 3 Bleo). Briefly 1.10^6^ cells were resuspended in 100µl of PBS, 2% BSA, 0.01% Tween and incubated with 10µl Fc Blocking reagent for 10 min 4°C then stained with 0.5µg of cell hashing antibody for 20 minutes at 4°C. After washing with PBS, 2% BSA, 0.01% Tween samples were counted and merged at the same proportion, spun 5 minutes 350 x g at 4°C, resuspended in PBS supplemented with 0.04% of bovine serum albumin at final concentration of 500 cells/µl and pooled sample were immediately loaded on Chromium.

#### Generation of single cell librairies, 10X Genomics scRNA-seq and data processing

Single-cell capture was performed using the 10X Genomics Chromium device (3′ V3). Single-cell libraries were sequenced on the Illumina NextSeq 500. Alignment of reads from the single cell RNA-seq libraries and unique molecular identifiers (UMIs) counting were performed with 10X Genomics Cell Ranger tool (v3.0.2). Reads of oligonucleotides tags (HTOs) used for Cell Hashing were counted with CITE-seq-Count (v1.4.2). Counts matrices of total RNA and HTOs were thus obtained for the 6 sequencing run, respectively named YoungD14, YoungD28, YoungD60, OldD14, OldD28 and OldD60.

#### Single cell data secondary analysis

All downstream analyses were carried out with Seurat R package (v4.1.0).

##### HTOs demultiplexing

For each 6 sequencing run, RNA and HTOs counts matrices were integrated into a Seurat object. HTOs counts were demultiplexed with HTODemux function to assign mice-of-origin for each cell (3 PBS-treated and 3 Bleomycin-treated). Only the cells identified as “Singlet” and passing quality control metrics (more than 200 detected genes, less than 0.25% of mitochondrial content) were kept for each Seurat object. The 6 seurat objects containing the raw counts of filtered cells and their metadata (age, day of lung collection post injection, treatment, mice-of-origin) were merged.

##### Samples integration

The computational integration of the different samples was performed using reciprocal PCA, as described in the dedicated vignette from Satija lab website (https://satijalab.org/seurat/articles/integration_rpca.html). Briefly, a “detailed samples” metadata column was created by aggregating age, day of lung collection post injection and treatment (ex: Young.D14.Bleo, Young.D14.PBS, etc). According to this “detailed samples” metadata, cells were split into 12 seurat objects. For each of 12 datasets, raw counts were normalized using SCTransform with Gamma-Poisson generalized linear model to estimate regression parameters. 3000 features were selected for being repeatedly variable across the 12 datasets. The sctransform residuals corresponding to the selected 3000 features were computed if they were missing in a particular object. Principal Component Analysis (PCA) was run individually for each object on the sctransform normalized counts of the 3000 selected features. A set of common anchors was found among the selected features, in dimensionally reduced data by reciprocal PCA, using FindIntegrationAnchors function with the following arguments: normalization.method = “SCT”, dims = 1:50, k.anchor = 5. This set of anchors was used to create a combined object containing integrated data using 50 dimensions for the anchor weighting procedure.

##### Clustering and annotation

First, PCA was run on the integrated data. UMAP and kNN clustering were computed on the first 80 computed principal components (PCs). Clusters corresponding to aggregates of cells sharing a low-UMIs content were discarded. The entire integration and clustering processes were rerun on the cleared dataset, following the exact steps described above. The finally obtained clusters were annotated on the basis of the expression of their specific markers listed in the output of the FindAllMarkers function. Cell type–specific marker genes were established using the Wilcoxon rank sum test with P values adjusted for multiple comparisons using the Bonferroni method. Adjusted P values <0.05 were considered significant.

*Differential expression analyses* between fibrotic and PBS conditions in all cell populations. Differential expression analyses between cells from bleomycin and PBS treated mice were carried out with DESeq2. First, all counts of each cell population from the same mice were aggregated to create a pseudobulk sample. For the PBS condition, we thus systematically obtain 18 pseudobulks samples corresponding to the 3 young and 3 old PBS-treated lungs collected at day 14, at day 28 and at day 60. For the fibrotic condition, we excluded the samples from bleomycin-treated lungs collected at day 60, which correspond to an almost complete fibrosis resolution. We thus obtained 12 pseudobulks corresponding to the 3 young and 3 old bleomycin-treated lungs collected at day 14 and at day 28. For the alveolar macrophages pseudobulks, all counts from the 3 subpopulations (AM1, AM2, AM3) were aggregated. For gCap and aCap pseudobulks, all counts of gCap plus Lrg1^pos^ gCap or aCap plus Lrg1^pos^ aCap were aggregated. Genes considered as differentially expressed were selected according to an adjusted p-value threshold of 0.05, obtained with the Wald test and the Benjamini-Hochberg method for multiple tests correction implemented in DESeq2.

#### Detailed analysis of endothelial cells (EC)

##### Identification of EC subpopulations

The clusters corresponding to general capillary cells (gCap) and aerocytes (aCap) were extracted and subclustered using the 80 PCs already computed from the integrated assay of the complete dataset. Clusters corresponding to Lrg1^pos^ aCap, Lrg1^pos^ gCap and sCap subpopulations were added to the annotation and UMAP was rerun on the subset of cells corresponding to gCap, aCap, proliferating EC, Lrg1^pos^ aCap, Lrg1^pos^ gCap, sCap, PV EC and Arterial EC using the same number of dimensions. The original raw counts were finally log-normalized with NormalizeData for all data exploration and visualization.

##### Relative subpopulations frequencies

All PBS-treated mice with the same age and sacrificed at D14, D28 or D60 were considered as replicates (n=9), while Bleomycin-treated mice were separated according to age and time of sacrifice (n=3 for each time point). The relative subpopulations frequencies at the 4 time points (PBS, D14, D28, D60) were obtained by calculating the ratio between the number of cells in each subpopulation and the total number of cells in each mouse replicate, either mixing or distinguishing young and old mice. Differential abundance analyses were carried out using edgeR (v3.32.1) to model overdispersed numbers of cells per group with the NB GLM method and then to test for differences in abundance between sample groups with the glmQLFTest function, which performs empirical Bayes quasi-likelihood F-tests and FDR controlling.

##### Bleomycin-induced fibrosis score

To compute a bleomycin-induced fibrosis score in gCap, aCap and PVEC, only the most significant upregulated genes from the differential expression analyses described above were selected according to adjusted p-value (<0.05), log2FoldChange (>0.5) and base mean expression (>10 for gCap, >5 for aCap and PV EC). Lists of selected genes were used to compute a score at the single cell level using the AddModuleScore function. Scores were finally compared between the 4 time points (PBS, D14, D28, D60) and between young or old animals, using the Wilcoxon rank sum test implemented in the FindMarker function.

##### Comparison between old vs young gCap in fibrotic and physiological conditions

To compare the transcriptomic profiles of gCap from old and young mice in fibrotic condition, counts from bleomycin-treated gCap and Lrg1^pos^gCap were aggregated within the same mouse for day 14 and day 28 time points, in order to obtain 6 young and 6 old pseudobulks samples. The same process has been applied to build pseudobulk samples of physiological gCap but including day 60 samples, resulting in 9 pseudobulks for either young of old groups. Differential analysis between old and young pseudobulks samples were carried out with DESeq2. DEGs were selected according to adjusted p-value (<0.05), absolute log2FoldChange (>0.5) and base mean expression (>5).

##### Systemic EC transcriptomic signature in fibrotic condition

As systemic EC from control samples were poorly detected either in mouse or human data, systemic EC from fibrotic conditions were compared to control systemic EC plus control PV EC using the pseudobulk-based approach.

##### Ingenuity Pathway Analysis

To investigate the enriched pathways and functions associated to DEGs either in the fibrotic versus control comparison or old versus young comparison, we used the core analysis of the Ingenuity Pathway Analysis tool (Ingenuity® Systems, www.ingenuity.com) using the results of the treatment (bleomycin)- or aging-perturbed genes obtained by DGE testing (see above) as input.

#### RNA velocity analyses

The RNA velocity analyses were carried out following the kallisto | bus (https://bustools.github.io/BUS_notebooks_R/velocity.html) and scVelo workflows (https://scvelo.readthedocs.io/).

##### Generation of spliced and unspliced matrices

Intronic sequences of the mouse genome assembly GRCm38 were first identified using the get_velocity_files function of the BUSpaRse R package (v1.5.2) and the gencode annotation (v.M25). In order to allow the pseudoalignment of reads on both cDNA and identified intronic sequences, indexes for both cDNA and intronic sequences were built using kallisto (v0.46.2). The spliced and unpliced matrices were then generated for each sequencing run using the wrapper combining kallisto and bustools (v0.39.2) functions, typically with the following command line: kb count -i mm_cDNA_introns_index.idx -g tr2g.tsv -x 10xv3 -o kb \ -c1 cDNA_tx_to_capture.txt -c2 introns_tx_to_capture.txt --workflow lamanno \ sample_x_L001_R1_001.fastq.gz sample_x_L001_R2_001.fastq.gz \ sample_x_L002_R1_001.fastq.gz sample_x_L002_R2_001.fastq.gz \ sample_x_L003_R1_001.fastq.gz sample_x_L003_R2_001.fastq.gz \ sample_x_L004_R1_001.fastq.gz sample_x_L004_R2_001.fastq.gz

##### Preprocessing

Generated spliced and unspliced matrices of each sequencing run were then preprocessed to filtered out empty droplets, to remove undetected genes and to replace ensembl IDs by gene symbols. Spliced and unspliced matrices corresponding to barcodes of subset of cells of interest, for example sCap EC, Lrg1^pos^ gCap and gCap from Bleomycin and PBS-treated mice at day 14, were integrated in a seurat object. The spliced raw counts were normalized using SCTransform from seurat R package, and PCA was run on these normalized data, as well as UMAP with the first 30 PCs. The seurat object was then converted to .h5ad file, keeping only the two raw spliced and unspliced assays.

##### RNA velocity using scVelo

The .h5ad file was read to an AnnData object using scVelo python package (v0.2.4). Again, data were preprocessed: the counts of the 3000 most variable genes with at list 20 reads were log-normalized using the filter_and_normalize function. PCA and nearest neighbours were computed before the estimation of velocity. The full transcriptional dynamics of splicing kinetics was resolved using the generalized dynamical model and represented as latent time in each cell.

#### NicheNet analysis

We used the nichenet R package (v1.0.0) ^37^ to infer ligand-receptor interaction candidates able to induce the expression of pro-angiogenic genes in sCap, Lrg1^pos^ gCap and Lrg1^pos^ aCap cells. For each of these 3 subpopulations, alternatively considered as receiver cells, we computed the activities of ligand sent by all the detected populations of the whole dataset. As genesets, we alternatively used the pro-angiogenic genes expressed in at least one third of cells with an average expression above 1 in the particular subpopulation. For each ligand activities computed for each subpopulation, we selected the top 14 most likely to induce the geneset of interest by ranking them according to Pearson’s correlation coefficient. The union of the 3 lists of prioritized ligands resulted in the 17 ligands used for the visualization. The selected receptors of those 17 ligands were only those considered as “*bona fide*”, i.e. those whose interaction with their ligand is not only predicted *in silico*. The only exception is for Apoe which didn’t have a *bona fide* receptor according to the NicheNet model, for which its best predicted receptor Scarb1 was added. The circos plot linking prioritized ligands and their receptors was realized following the steps described in the dedicated nichenet’s vignette (https://github.com/saeyslab/nichenetr/blob/master/vignettes/circos.md).

#### RNA-FISH

RNAscope probes against mmu-Lrg1 (ID: 423381), mmu-Aplnr (ID: 436171),mmu-Ednrb (ID: 473801), mmu-Col15a1 (1092391), mmu-Hif1a1 (313821), mmu-Cd74 (437501), mmu-Sox17 (433151), mmu-Vegfa (405131) were prepared and obtained from Advance Cell Diagnostics. FISH assays were performed on FFPE lung sections, as previously described [1] using Multiplex Fluorescent Reagent Kit V2 (Advanced cell Diagnostics) and TSA Plus Cyanine 3 and Cyanine 5 (Perkin Elmer) as fluorophores according to manufacturer’s recommendations. Acquisition was performed using an inverted confocal microscope LSM 710 (Zeiss) and a 40x oil immersion lens.

### Immunohistochemistry and Immunofluorescence microscopy

Five-µm paraffin-embedded sections were deparaffinized/rehydrated and H&E stained as previously described^56^. After washing, the sections were antigen-retrieved using citrate buffer (pH 6.0; Sigma-Aldrich). Autofluorescence was quenched for 20-30 minutes in NH4Cl buffer (0.1M) and passed in Photobleacher for 1h30. Slides were incubated with the primary antibodies (anti-CD31 (1:50), anti-COL15A1 (1:100), Anti-SMA–FITC (1:500)) diluted in a blocking solution BSA (1%) over night at 4°C. After washing in PBS-T (0.05%), slides were incubated for 1h with secondary antibodies (1:400) at room temperature. Nuclei were counterstained with DAPI (1:5000, D1306 Life Technologies). Slides were mounted using Immu-Mount (9990414 Fisher scientific). Representative regions of stained slices were digitalized on an epifluorescent Zeiss microscope and analyzed using the software FIJI.

### Western blot

Cells were lysed with RIPA cell lysis buffer (Thermofisher) containing phosphatase and protease inhibitors (Invitrogen). Proteins were separated on commercial gradient (4-12%) acrylamide gels and transferred onto PVDF membranes. Proteins were detected by overnight incubation at 4°C with the indicated primary antibodies followed by 1h incubation at room temperature with horseradish peroxidase-coupled secondary antibody and enhanced chemiluminescent substrate (PerkinElmer, Waltham, USA) using a LasImager (GE HealthCare, Chicago, Illinois, USA). Blots were run on parallel gels when proteins of similar molecular weight were analyzed. The western blot quantification was performed with the help of the ImageJ software.

### Taqman Real time PCR

Total HMEC1 RNA was isolated using trizol (Thermo Fisher Scientific) according to the manufacturer’s instructions. RNAs (1 µg) were reverse-transcribed with High capacity cDNA reverse transcription kit (Thermo Fisher Scientific), and PCR was performed using Taqman Real time PCR technology (Applied Biosciences). All reactions were done using Real-Time PCR system (Thermo Fisher Scientific), and expression levels calculated using comparative CT method (2 DDCT). Gene expression levels were normalized to RPLP0 level. The following probes from Taqman (Applied Bioscience) were used: Hs-RPLP0 (99999902-m1), Hs-VEGFA (00900055-m1) and Hs-SMAD2 (00183425-m1).

### Re-analysis of public scRNA-seq datasets

Human lung fibrosis ^34^ and mouse lung public datasets ^35^ were re-analyzed using reciprocal PCA, in order to integrate samples from different patients or different mice. To reduce the complexity of the human lung fibrosis dataset, we only kept cells coming from either IPF-diagnosed or normal samples and we randomly sampled 10000 immune cells and 5000 epithelial cells to reduce the dataset dimensionality. Adding the total number of endothelial and mesenchymal cells, we obtained a dataset of 25816 cells. Differential expression and differential abundance analyses were carried out on this dataset in the same way as for our bleomycin dataset as described above.

### Statistical Analysis

Statistical analyses were performed using GraphPad Prism and R studio (v.4.1.2). Boxplots are given as median ±quartile. Histogram and scatter plot are given as mean ±SD. Two-tailed Mann–Whitney test or multiple paired t test were used for single comparisons; Two-way ANOVA test followed by a multiple comparisons test with Holm-Šídák correction was used for multiple comparisons. P-value <0.05 was considered statistically significant.

## DATA AVAILABILITY

The scRNA-seq and spatial transcriptomic data sets have been deposited in the Gene Expression Omnibus SuperSerie GSE234199.

## CODE AVAILABILITY

The scripts used for the analysis of scRNA-seq and spatial transcriptomic data are available on github : https://github.com/marintruchi/Aging_affects_reprogramming_of_PCEC.

## Acknowledgements

The authors thank the technical support of the UCA GenomiX, the CoBioDA bioinformatics hub of the IPMC and the microscopy facility from the IPMC, part of the « Microscopie Imagerie Cytométrie Azur» GIS IBiSA labeled platform. We also thank the staffs from the Nice Hospital-Integrated Biobank (BB-0033-00025) and the Giessen PNEUMObank as well as from the animal care facilities institutions at Sophia Antipolis (IPMC Animal Care Facility) and Lille (High Technology Animal Care Facility, University of Lille 2). We are also grateful to A. Monteil and C. Lemmers from the Vectorology facility, PVM, Biocampus Montpellier, CNRS UMS3426. We acknowledge the support from the Centre National de la Recherche Scientifique (CNRS), Institut National de la Santé et de la Recherche Médicale (Inserm), Université Côte d’Azur, Université de Lille, the French Government, through the National Research Agency (ANR) programs ANR-PRCI-18-CE92-0009-01 FIBROMIR / ANR-22-CE17-0046-01 MIR-ASO and the UCAJ.E.D.I. Investments in the Future project with the reference number ANR-15-IDEX-01, the Programme Contrat Plan État Région (CPER) – CTRL (Centre Transdisciplinaire de Recherche sur la Longévité, projet FISSURE) as well as Canceropôle PACA.

## Author contributions

BM, NP, GV and CC conceived and designed the study and supervised the entire work. MT and BM wrote the manuscript. GS, MGI, HC and AB performed in vivo studies. GS, MGI, HC, JF, NB and CdS performed biochemical and cellular biology experiments. GS, CdS and MJA performed the Visium spatial transcriptomic experiment. VM and CGR generated scRNA-seq data and primary analysis. MT performed bioinformatic analysis of scRNA-seq and spatial transcriptomic data. KLB co-supervised the computational analysis. GS, MGI, RL and HC performed and analyzed immunostainings. MT, GS, AL, NR, GV, CC, SB, NP and BM performed biological interpretation. VH, PH, AG, CHM and SL provided IPF-derived biological materials and contributed to biological interpretation of data. RR, MP, NR, PB and SB gave conceptual advices. OP and SB provided ressources. All authors read and corrected the final manuscript.

## Supplementary information

### Correspondence of supplementary tables

**Supplementary table S1.** Differential expression analysis between topics deconvoluted from spatial transcriptomic data.

**Supplementary table S2.** Differential expression analysis (Seurat) between annotated subpopulations of single cell RNA-seq data.

**Supplementary table S3.** Differential expression analysis (DESeq2) between lung cell types from bleomycin-treated and PBS-treated mice.

**Supplementary table S4.** Differential expression analysis (DESeq2) between IPF patients and healthy control for aCap, gCap, systemic EC and PV EC, from the Habermann et al. Dataset^34^.

**Supplementary table S5.** Differential expression analysis (DESeq2) between old and young aCap, gCap and PVEC cells in fibrotic condition.

**Supplementary table S6.** Differential expression analysis (DESeq2) between old and young aCap, gCap and PVEC cells in physiological condition.

### Correspondence of supplemental Figures

**Supplemental Figure S1. Histological and spatial transcriptomics data integration of injured lung slices from young and old mice.**

**Supplemental Figure S2. scRNA-seq analysis of young and aged mouse lungs following BLM challenge at 3 time points.**

**Supplemental Figure S3. Lrg1 expression in lung cell populations.**

**Supplemental Figure S4. LRG1 affects the TGF-β signalling towards a pro-angiogenic response.**

**Supplemental Figure S5. Bleomycin-induced PCEC subpopulations are associated with pro-angiogenic signaling.**

**Supplemental Figure S6. Expression of bleomycin-associated signatures in PCEC from young and aged mouse lungs.**

**Supplemental Figure S7. Expression of selected genes according to RNA Velocity latent time analysis of PVEC subpopulations.**

