## Supplemental Figures for "Aging affects reprogramming of murine pulmonary capillary endothelial cells after lung injury"

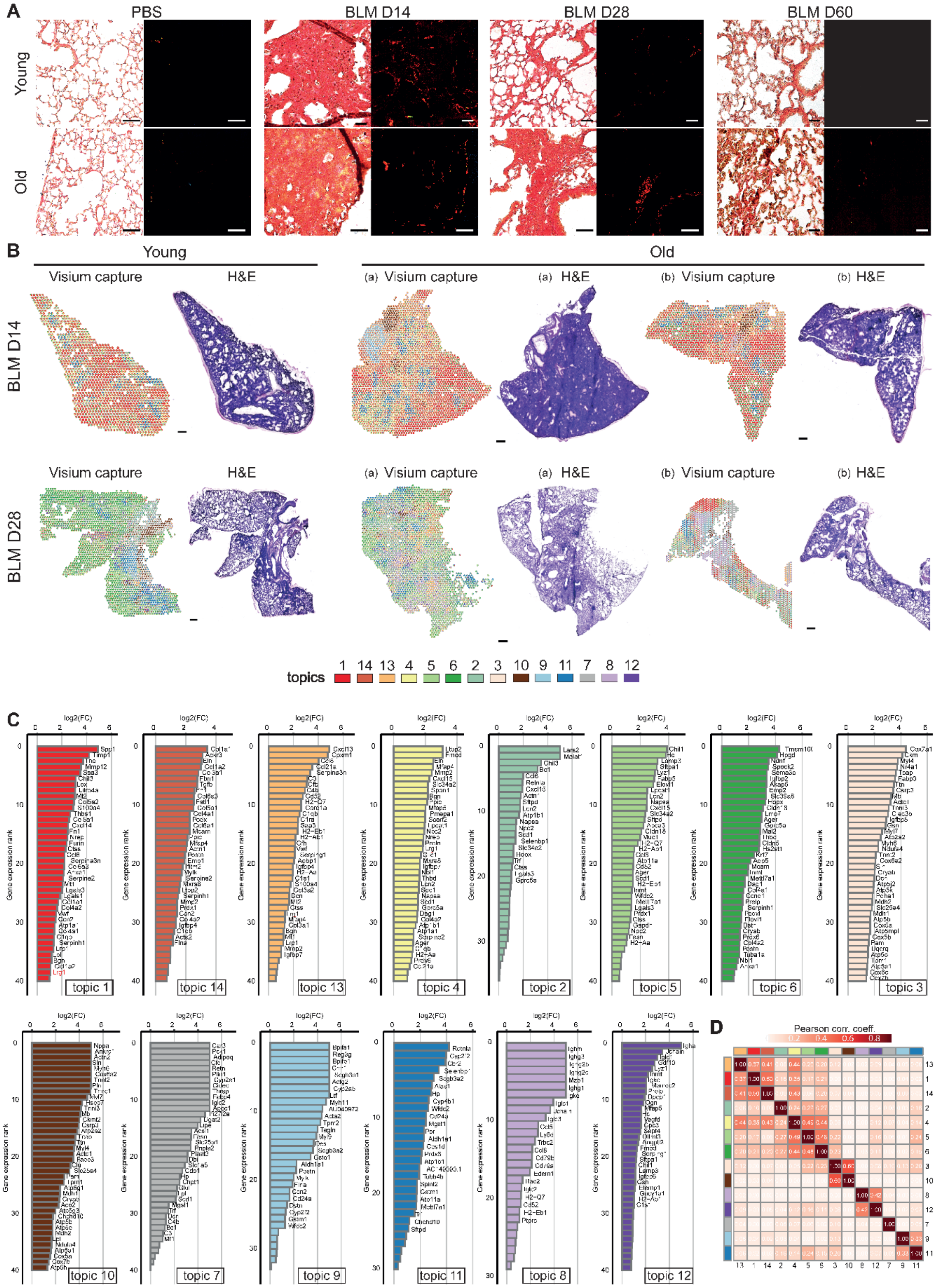


**Supplemental Figure S1. Histological and spatial transcriptomics data integration of injured lung slices from young and old mice. (A)** Representative image of picrosirius red staining of collagen deposition on lung parenchyma from PBS or bleomycin (BLM)-treated young or old mice, 14, 28 or 60 days after treatment under phase-contrast (left) or polarized light (right) microscopy. Scale bars=100µm. **(B)** Spatial transcriptomics data from histological sections (H&E) of lungs from young (n=1) and old (n=2) mice challenged with bleomycin and collected at fibrotic peak (day 14) and during regeneration (day 28). Spots are colored on the Visium capture according to assigned topics. Scale bars=200µm. **(C)** Top genes markers of each topics. **(D)** Heatmap of topics correlation based on gene expression.

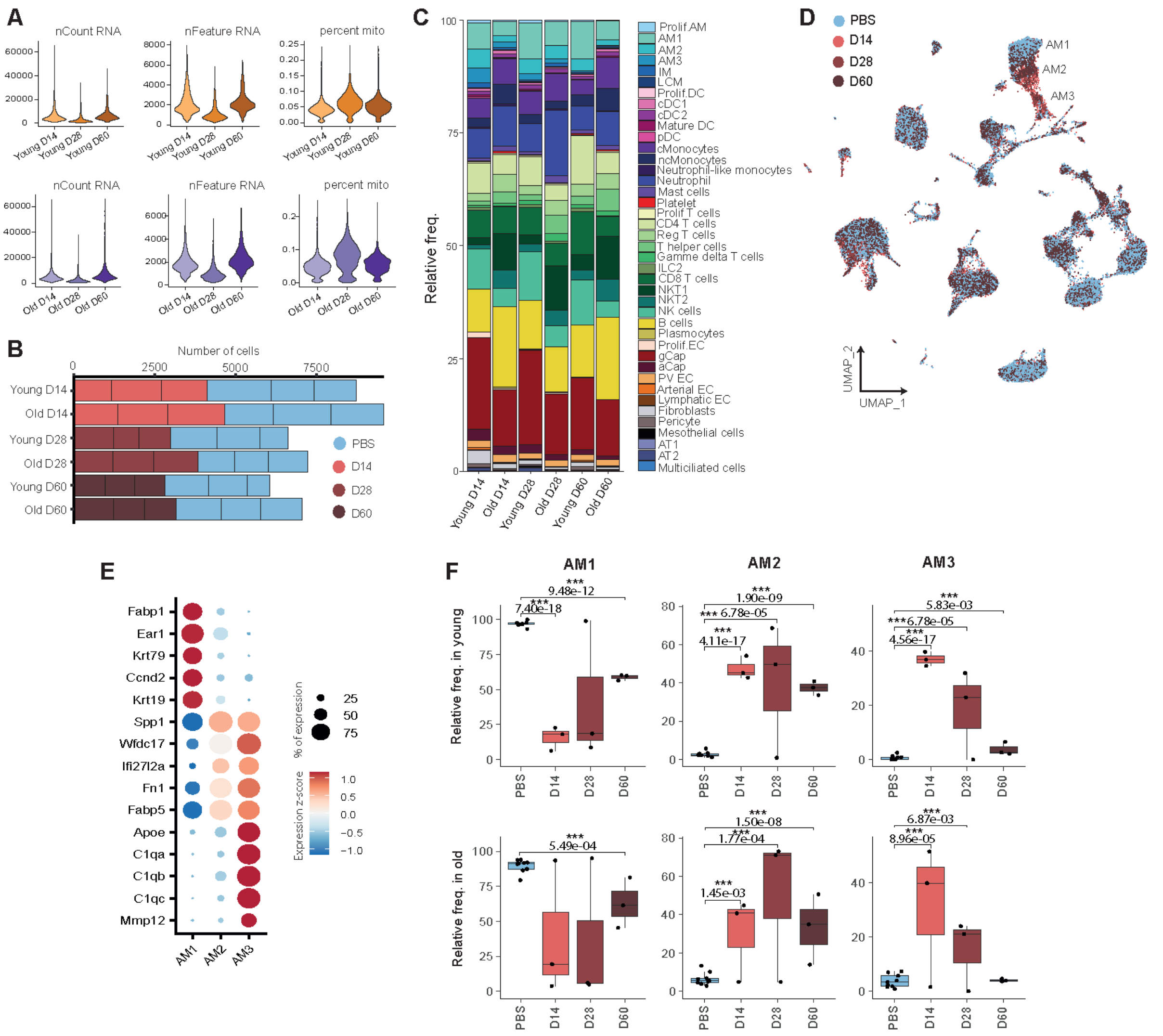

**Supplemental Figure S2. scRNA-seq analysis of young and aged mouse lungs following BLM challenge at 3 time points.** (**A**) Number of UMI, detected genes and percentage of mitochondrial content for each sequenced time point in young (top) and old (bottom) mice. (B) Number of cells by mice grouped by conditions, timepoints and age. (**C**) Relative proportions of all populations defined in integrated dataset. (D) UMAP of the integrated dataset. Cells are colored according to the timepoint. AM= alveolar macrophages. (E) Top genes markers in AM subpopulations (AM1, AM2 and AM3). (F) Relative proportions of alveolar macrophages subpopulations between young and old mice across time points.

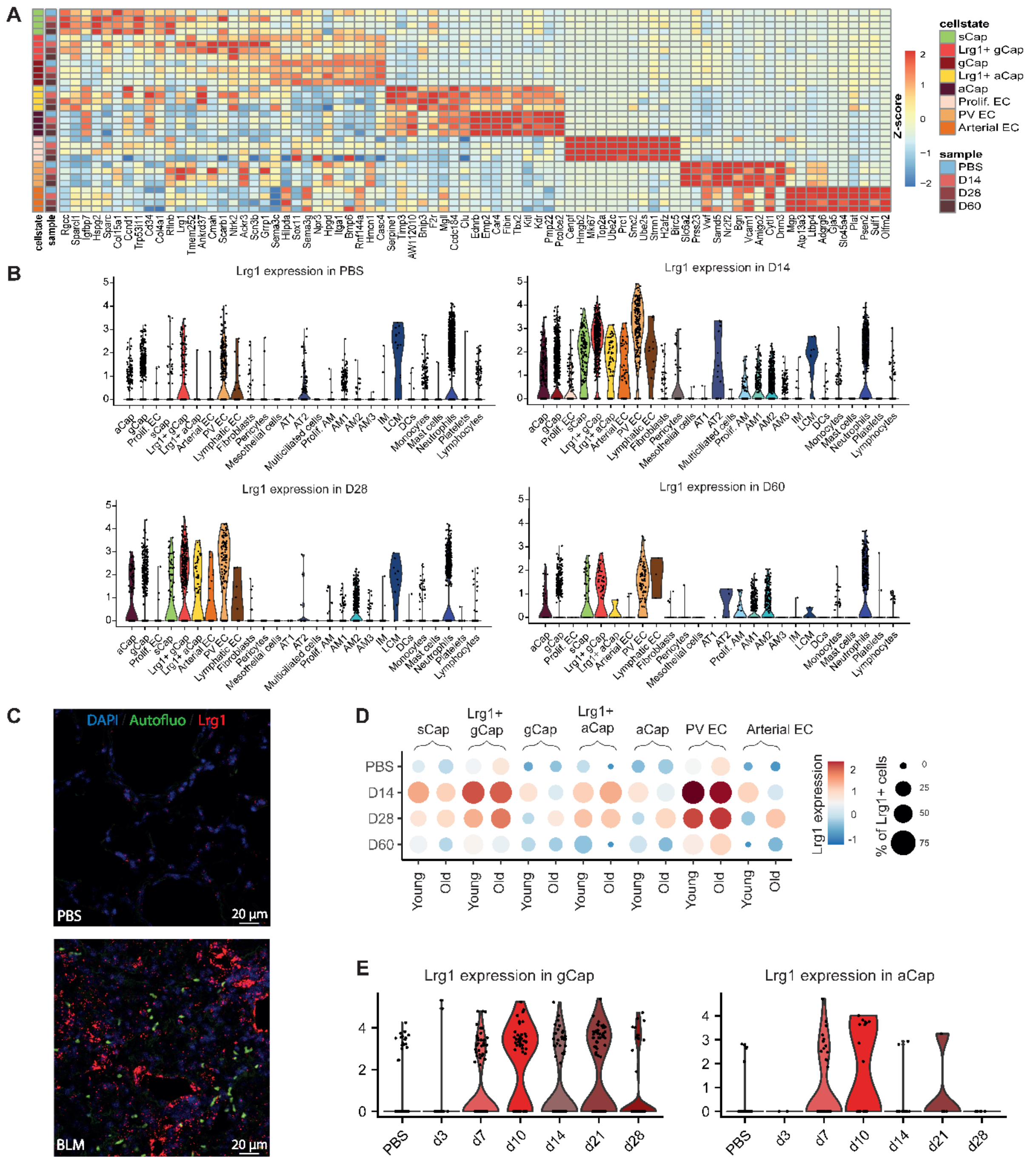

**Supplemental Figure S3. Lrg1 expression in lung cell populations. (A)** Heatmap of top EC subpopulations markers. All cells from young and old mice from the same condition are pooled. Expression is indicated as a z-score. (**B)** Normalized expression of Lrg1 in all cell populations in PBS-treated mice and in bleomycin-treated mice at indicated time points. (**C)** *In situ* hybridization of Lrg1 mRNA in PBS-treated (up) or Bleomycin-treated (down) lungs of young mice at D14. Representative image (n=3). Nuclei are counterstained with DAPI. (**D)** Normalized expression of Lrg1 comparison between pulmonary endothelial subpopulations and between young and old animals at indicated time points. (**E)** Normalized expression of Lrg1 in gCap and aCap isolated from Strunz et al. dataset (35).

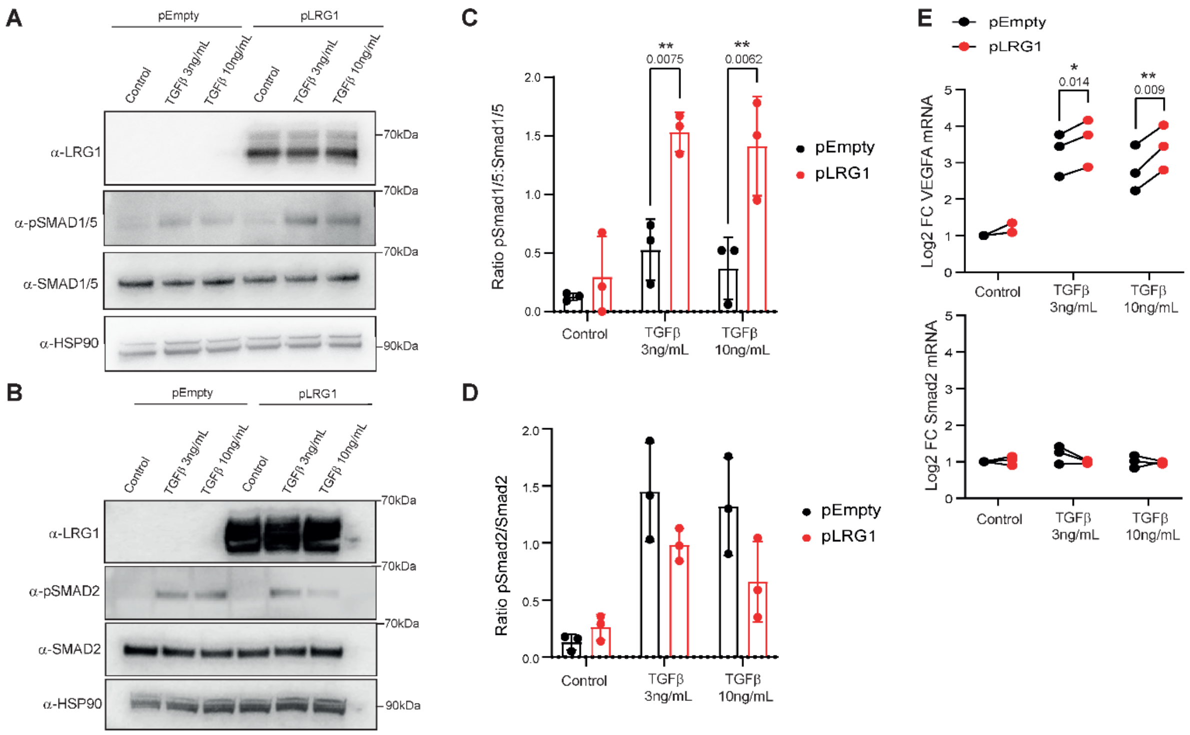


**Supplemental Figure S4. LRG1 affects the TGF-β signaling towards a pro-angiogenic reponse.** HMEC1 human endothelial cell line were transduced with an empty or LRG1 (pLRG1) lentiviral particles. Transducted cells were stimulated 1 hour with TGF-β1 at 3 or 10 ng/mL or not (Control). ***(A-B)*** Western blot of pSmad1/5 and total Smad1/5 (**A**) or pSmad2 and total Smad2 (**B**) on protein lysates from pEmpty or pLRG1 cells under TGF-β1 stimulation. LRG1 and HSP90 protein level were revealed to validate the transduction efficacy and the loading, respectively. Experiments were conducted at least 3 times. (**C-D**) Quantification of Phosphoprotein/total protein ratio for Smad1/5 (**C**) or Smad2 (**D**) in different experiments. Statistical analysis: ANOVA Two way with pairwise comparison (Bonferroni correction) (**E**) RT-qPCR of VEGFA and SMAD2 mRNA from pEmpty or pLRG1 cells under TGF-β1 stimulation. Log2 fold change were normalized by RPLP0 expression and on pEmty control conditions. Experiments were conducted at least 3 times. Statistical analyses: Multiple paired t-test.

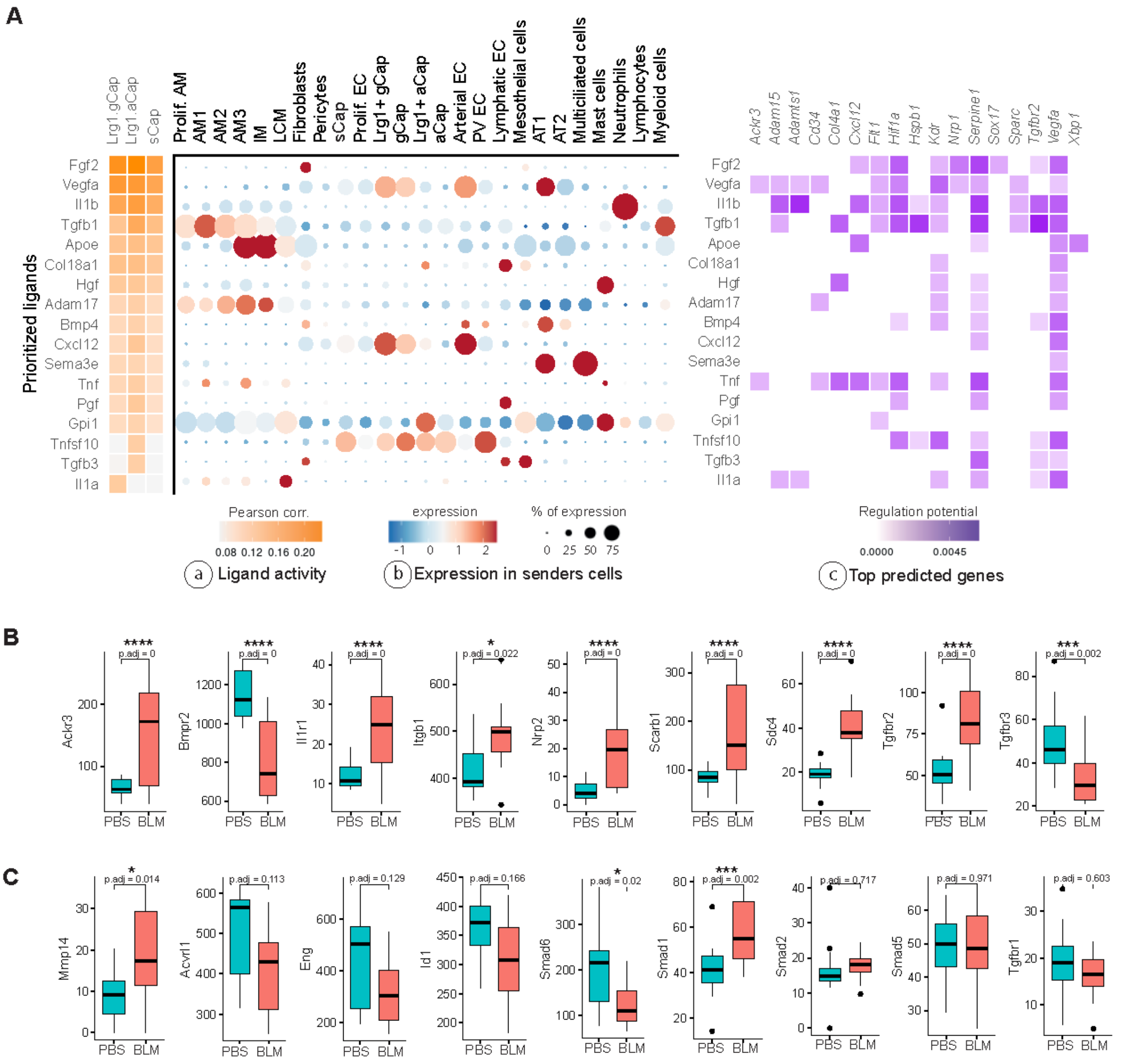


**Supplemental Figure S5. Bleomycin-induced PCEC subpopulations are associated with pro-angiogenic signalling. (A)** NicheNet analysis inferring upstream ligands most likely to induce a pro-angiogenic signature in Lrg1$^{pos}$ EC subpopulations: (a) Prioritized ligands are ranked according to their potential activity; (b) Level and percentage of expression of the prioritized ligands; (c) Top predicted target genes of prioritized ligands. **(B)** Differentially-expressed receptors between bleomycin and PBS treated gCap. **(C)** Expression comparison between bleomycin and PBS treated gCap for *Mmp14*, Bmp9-associated signaling genes *Acvrl1, Eng, Id1* and *Smad6*, as well as *Smad1*, *Smad2*, *Smad5* and *Tgfbr1*.

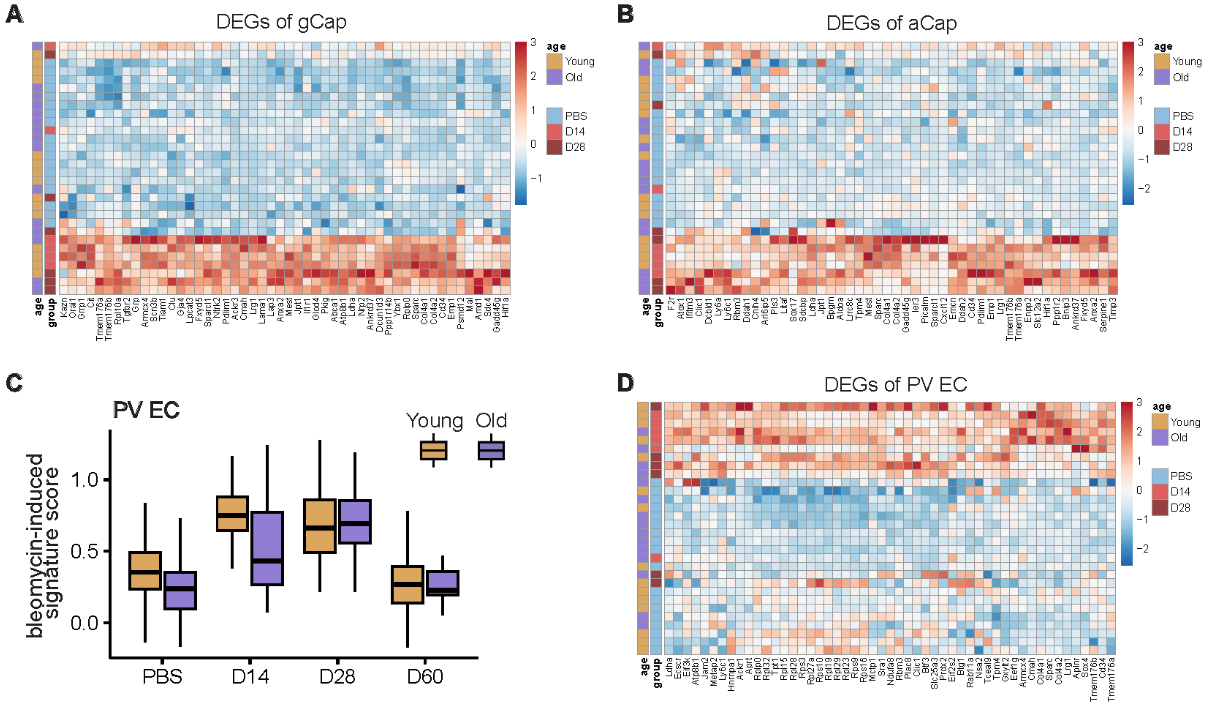


**Supplemental Figure S6. Expression of bleomycin-associated signatures in PCEC from young and aged mouse lungs.** **(A)** Heatmap of top 50 DEGs between Bleomycin and PBS treated gCap. **(B)** Heatmap of top 50 DEGs between Bleomycin and PBS treated aCap. **(C)** Bleomycin (BLM)-induced signature score (see material and methods section) in PV EC in young and old mice across time points. **(D)** Heatmap of physiological and pathological PV EC markers in spatial transcriptomics data.

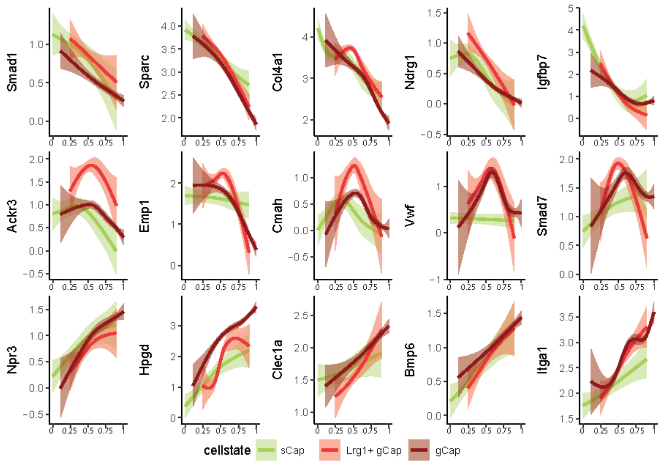


**Supplemental Figure S7. Expression of selected genes according to RNA Velocity latent time analysis of PVEC subpopulations** (sCap, Lrg1$^{pos}$ gCap and gCap*).* The cell population (cellstate: sCap, Lrg1$^{pos}$ gCap and gCap) are indicated with different colors.
